# Transmembrane PhoxID: Photo-proximity labelling across the plasma membrane

**DOI:** 10.64898/2026.01.15.699634

**Authors:** Fátima Yuri Tanimura Valor, Mikiko Takato, Ayane Araki, Seiji Sakamoto, Tomonori Tamura, Itaru Hamachi

## Abstract

Transmembrane proteins perform essential roles in cellular transport, signalling, and communication. The function and dynamics of these proteins are precisely regulated by interactions on both sides of the plasma membrane; thus, mapping the composition of these interactomes is a fundamental challenge in molecular biology. Proximity labelling methods are powerful tools for this purpose; however, existing approaches that rely on membrane-impermeable reactive species are limited by the ability to detect only one side, either the extracellular or the intracellular region, of transmembrane proteins. Here, we capitalized on the cell permeability of singlet oxygen to carry out proximity labelling of the cytoplasmic side of transmembrane proteins using an extracellularly anchored photosensitizer. We applied this method, termed transmembrane PhoxID (tmPhoxID), to several receptors (GRID2, GABA_A_R, and GRM1) in the living mouse brain and successfully determined their specific intracellular interactomes. Notably, network analysis of the identified proteins revealed that this method can characterize the native components of transsynaptic nanocolumns formed at parallel fibre–Purkinje cell synapses. Furthermore, our study revealed a previously uncharacterized GABA_A_R-CAMKV interaction in mice and human brains. Our results provide a proof of concept for transmembrane and transcellular proximity labelling, providing a powerful platform for analysing the interactome of transmembrane proteins.

## Introduction

Transmembrane proteins (TMPs) are integral membrane proteins that span both the extracellular and intracellular environments and function as receptors, transporters, channels, enzymes, and structural proteins, enabling cells to respond rapidly to changes in their surrounding environment^1,2^. They are precisely regulated by an elaborate network of protein–protein interactions and form dynamic, unique multimolecular complexes with different characteristics on opposite sides of the plasma membrane^2,3^. Therefore, identifying the composition of the intracellular and extracellular interactomes, as well as how they change in response to environmental stimuli, is fundamental to elucidating the molecular machinery of TMPs. Traditional co-immunoprecipitation (Co-IP) coupled with mass spectrometry has long been the standard method for investigating the cellular protein networks associated with TMPs^4,5^. However, this method lacks the spatiotemporal resolution necessary to capture the localized and transient interactions that are expected for transmembrane signal transduction events. Additionally, the use of cell/tissue homogenates poses the problem of potential false positives and false negatives^6^. Proximity labelling (PL) provides a valuable means to sidestep these limitations and has been widely used to analyse the extracellular and intracellular proteomes of TMPs^7–19^. However, most of the reactive species generated by enzymes or photocatalysts for PL are membrane-impermeable or poorly membrane permeable, and labelling is therefore largely restricted to proteins on the same side as the catalyst. As a result, simultaneous mapping of both intracellular and extracellular interactomes of TMPs has remained challenging. Moreover, many TMPs, such as those whose *N*- and *C*-termini are both exposed on the extracellular side, lack a cytoplasmic domain that is amenable to enzyme fusion, making it difficult to perform PL on the intracellular side^1^. Although these challenges may be resolved if the reactive species generated on one side of the plasma membrane can traverse the lipid bilayer, the feasibility of PL across the membrane has not been thoroughly validated.

In this study, we report that PhoxID, which we recently developed^20,21^, provides a unique platform for achieving this sort of “transmembrane PL”, as it employs membrane-permeable singlet oxygen (^1^O_2_) as a reactive species for PL^22–25^ (**Fig. 1a**). The established workflow for transmembrane PhoxID (tmPhoxID) enabled the detection of intracellular interactomes of various TMPs with high spatial resolution in cultured cells, ex vivo (acute brain slices), and *in vivo* (the living mouse brain). The applicability of tmPhoxID was demonstrated by targeting the δ2 glutamate receptor (GRID2), a key organizer of excitatory synapses^26^. This analysis successfully identified the intracellular components of parallel fibre–Purkinje cell synapses as well as the extracellular interactomes. The spatial resolution of tmPhoxID was further assessed by targeting two additional receptors—γ-aminobutyric acid type A receptor (GABA_A_R)^27^ and metabotropic glutamate receptor 1 (GRM1)^28^—which are coexpressed but not colocalized with GRID2 in the cerebellum. This analysis revealed the intracellular interactomes exclusive to each receptor, underscoring the receptor specificity and spatial precision of tmPhoxID even within subsynaptic microenvironments. Furthermore, tmPhoxID targeted to GABA_A_R successfully identified the previously uncharacterized GABA_A_R–CAMKV interaction *in vivo*.

**Figure 1.**
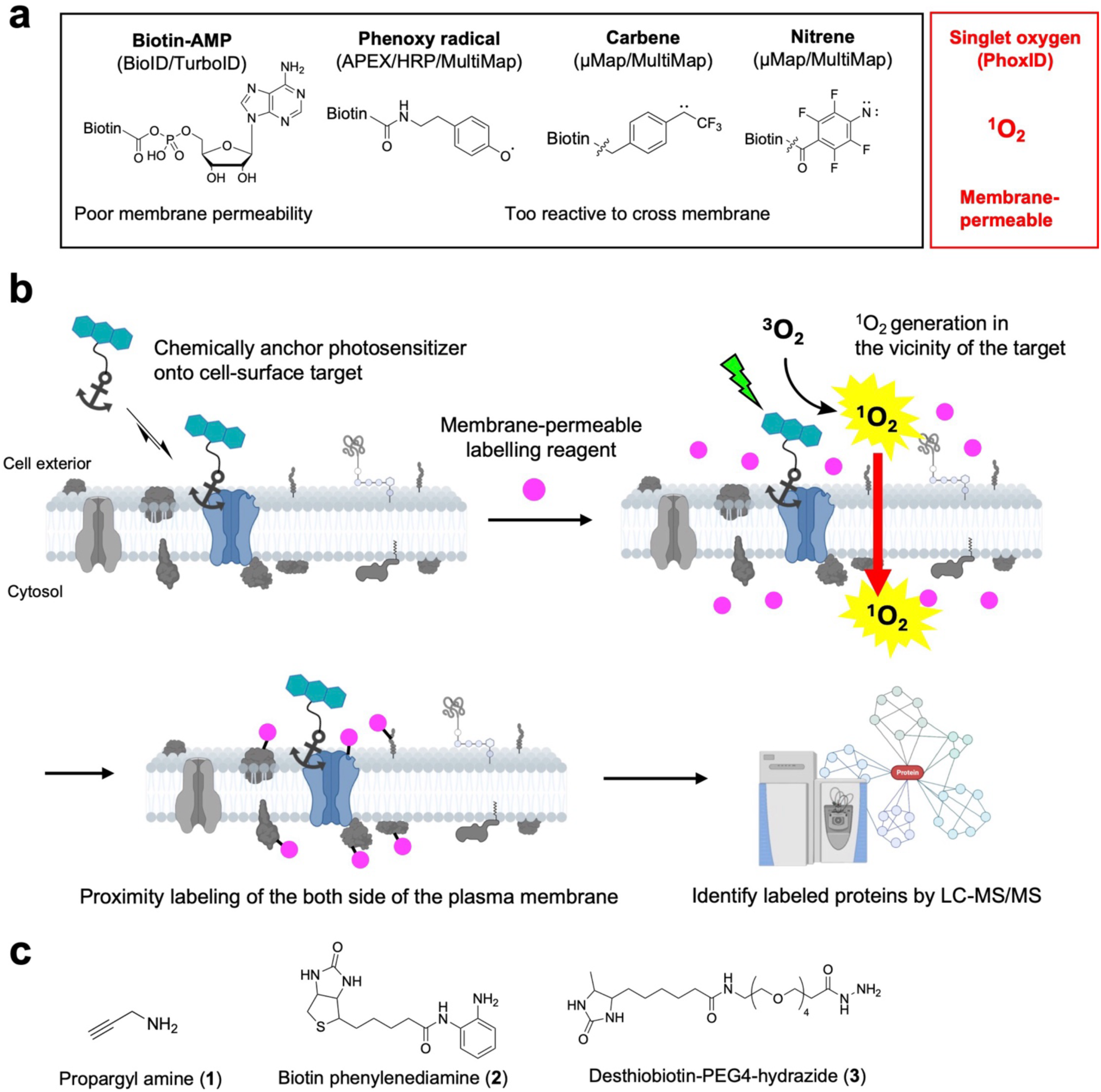
Concept of tmPhoxID. (a) Reactive species utilized in different PL strategies, highlighting those that are restricted to the extracellular space versus singlet oxygen, which can diffuse across membranes. (b) Schematic overview of the tmPhoxID workflow. First, the photocatalyst is anchored to the target protein. Upon the addition of a cell-permeable reagent and subsequent irradiation, the photocatalyst generates singlet oxygen. Owing to its ability to diffuse across the cell membrane, singlet oxygen enables the labelling of intracellular proteins, which can subsequently be identified and quantified by LC–MSMS. (c) Labelling reagents evaluated in the context of tmPhoxID.

## Results

### Proof-of-concept of tmPhoxID with cultured cells

On the basis of the labelling mechanism of PhoxID, we hypothesized that the ^1^O_2_ generated upon excitation of the photosensitizer could diffuse across the lipid bilayer and oxidize proteins on the opposite side of the membrane from the site where the photosensitizer is anchored (**Fig. 1b**)^20^. These oxidized proteins can subsequently be labelled with nucleophilic reagents bearing amine, aniline, or hydrazide groups to introduce tags for detection and enrichment (e.g., biotin, clickable moieties) (**Fig. 1c**). As a proof-of-concept, we aimed to demonstrate that a photosensitizer anchored to the intracellular side of the plasma membrane could mediate the labelling of proteins on the extracellular surface. This setup was previously used to demonstrate the cell impermeability of the reactive species generated in enzyme-based PL^8^. We expressed the HaloTag on the cytoplasmic side of the plasma membrane by fusing it to the inner leaflet protein Lyn. HeLa cells transfected with the Lyn-Halo were treated with cell-permeable acetylated dibromofluorescein-Halo ligand conjugate (**AcDBF-HL**, **4**) to covalently attach the DBF photosensitizer to the protein (**Extended Data Fig. 1a**). The DBF-anchored cells were then subjected to photoirradiation in the presence of **Hyd-PEG4-Bt** (**6**), and the labelled proteins were stained with cell-impermeable streptavidin-Alexa Fluor 647 (AF647) in live cells for the specific detection of cell-surface biotinylated proteins (**Fig. 2a, Extended Data Fig. 1c**). As we expected, cell-surface labelling was observed in cells expressing DBF-labelled Lyn-Halo upon photoirradiation (**Fig. 2b**). The biotin signal was absent in both the unirradiated samples and the cells lacking DBF anchoring, confirming that the labelling was driven by ^1^O_2_ produced by DBF (**Fig. 2b** and **Extended Data Fig. 1d**). These results suggest that ^1^O_2_ generated within the cells could diffuse across the lipid bilayer and mediate the labelling of proteins on the cell surface, validating the tmPhoxID concept.

**Figure 2.**
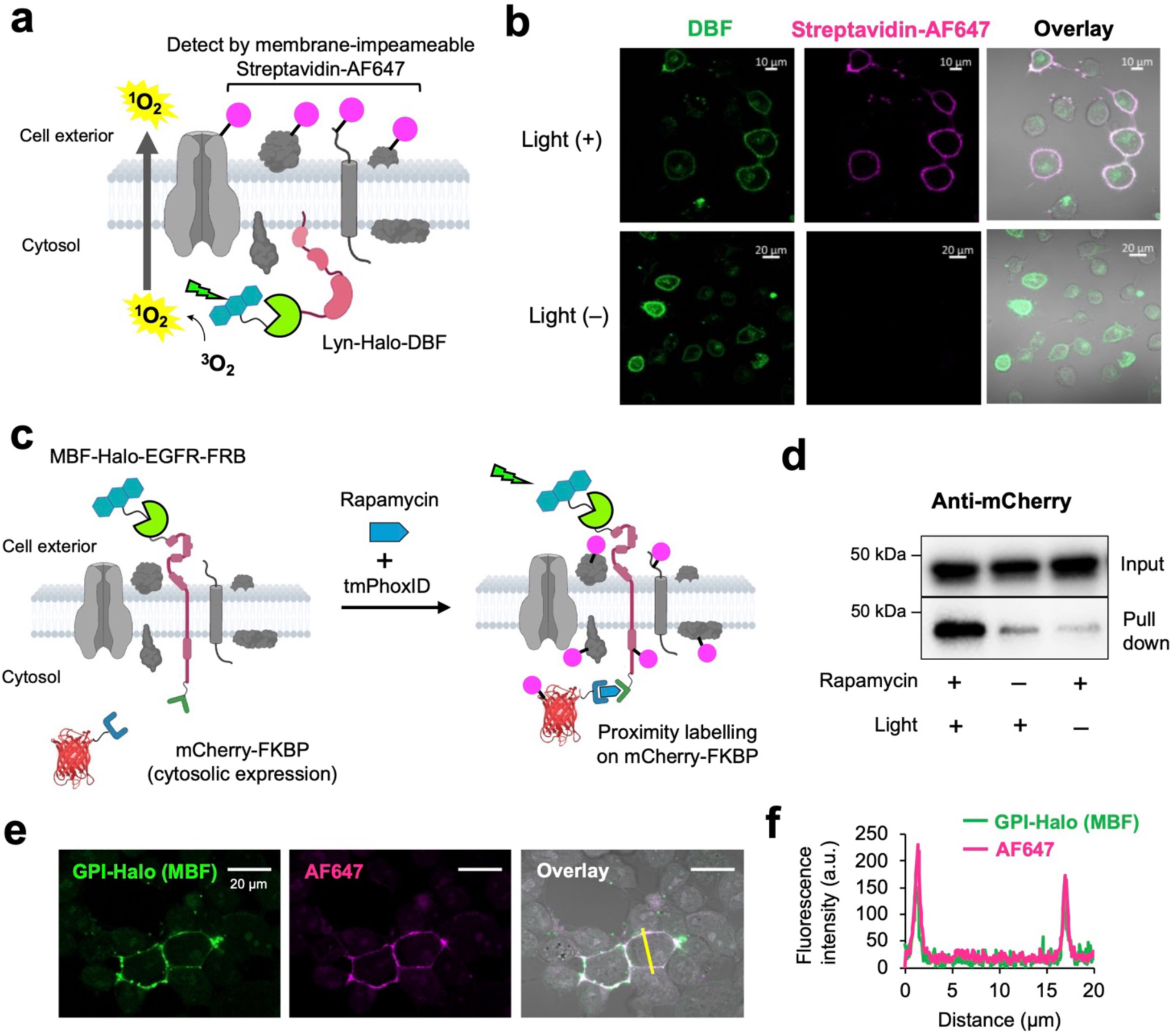
tmPhoxID can label proteins on the opposite side of the plasma membrane. **(a)** Schematic representation of the experimental design. First, the DBF photosensitizer was introduced to Lyn-Halo, which was localized on the inner leaflet of the plasma membrane. Subsequently, cells were illuminated in the presence of Hyd-PEG4-Bt, and biotinylated proteins on the cell surface were selectively visualized using Streptavidin-AF647. **(b)** CLSM images of biotinylated proteins on the HeLa cell surface. An intracellular photosensitizer facilitated the labelling of extracellular proteins. **(c)** Experimental design for the rapamycin-dependent labelling of cytosolic mCherry–FKBP12. **(d)** Pulldown detection of biotin-labelled mCherry–FKBP12. **(e)** CLSM images of HEK293T cells after tmPhoxID with MBF-modified GPI-Halo. Proteins labelled with **1** were visualized after cell fixation via a click reaction with AF647–N_3_. **(f)** Fluorescence intensity profile along the yellow line in panel **e**.

We next assessed whether this method could be used to monitor proteome dynamics at the inner leaflet side of the plasma membrane. As a test, we designed the model system shown in **Figure 2c**. HaloTag was fused to the N-terminus of epidermal growth factor receptor (EGFR) to anchor photosensitizer monobromofluorescein (MBF) to the extracellular side of the plasma membrane, and an FKBP-rapamycin-binding (FRB) domain was fused to the cytoplasmic C-terminus^29^. An mCherry-FKBP12 fusion protein was expressed in the cytosol. tmPhoxID was performed in the presence of propargylamine (**1**), which has been previously identified as an efficient cell-permeable labelling reagent for ^1^O_2_-mediated PL (**Fig. 1c**)^30^. In the basal state, the labelling degree of mCherry-FKBP12 was minimal, as this protein is not particularly enriched at the plasma membrane. However, upon treatment with rapamycin, the labelling signal was increased three- to four-fold compared with that in the untreated control, accompanied by the recruitment of mCherry-FKBP12 to the plasma membrane (**Fig. 2d**, **Extended Data Fig. 1e–g**). These data demonstrated that tmPhoxID can quantitatively track changes in the composition of the intracellular interactome of the target TMPs.

We further assessed the spatial resolution of tmPhoxID. Cells presenting MBF on their surface were irradiated in the presence of **1**, and the labelled proteins were subsequently visualized in the fixed cells by copper(I)-catalyzed azide-alkyne cycloaddition (CuAAC) with AF647-azide. Confocal imaging analysis revealed that PA labelling was restricted to the plasma membrane periphery (**Fig. 2e**, **f**), suggesting sub-micrometre spatial resolution and confirming that the labelling was limited to target-proximal proteins, even on the other side of the plasma membrane.

### GluD2-tmPhoxID at parallel fiber–Purkinje cell synapses ex vivo and *in vivo*

Having demonstrated the applicability of tmPhoxID in cell culture, we next investigated whether this method could identify the real membrane-spanning interactomes of endogenous TMPs *in vivo*. The delta-2 type glutamate receptor (GRID2) was selected as the initial target. GRID2 is a postsynaptic receptor that is highly enriched in cerebellar Purkinje cells, where it performs critical roles in synapse formation, plasticity, and motor coordination^26^. To determine the optimal labelling reagent for *in vivo* experiments, we evaluated compounds **1**–**3** (**Supplementary Fig. 1** for successful tmPhoxID with **2** and **3** in cultured cells). To selectively anchor the photocatalyst to the extracellular domain of endogenous GRID2, we injected the GRID2-specific nanobody–DBF (Nb^GRID2^–DBF) conjugate, developed in our previous study^20^, into the lateral ventricles (LVs) of the living C57BL/6N mouse brain (**Fig. 3a**). On the following day, the labelling reagent was injected into the cerebellum, and the same site was photo-irradiated through an optical fibre (520 nm) for 10 min. The labelled proteins in the cerebellum tissue were enriched, digested into peptides, and analysed by LC–MSMS in data-independent acquisition (DIA) mode (**Extended Data Fig. 2**)^31^. In photo-based PL, the light intensity is a key factor that determines the reaction radius^20,23^. Therefore, we first examined the light intensity dependence of tmPhoxID. Under the 0.16 W mm^-2^ condition used in conventional PhoxID, extracellular proteins were primarily detected, which was consistent with our previous reports. However, when the light intensity was increased to 0.96 W mm^-2^ or higher, the labelling of intracellular proteins was increased (**Extended Data Fig. 3, Supplementary Data 1**). On the basis of these observations, we set the light intensity to 0.96 W mm^-2^ for *in vivo* tmPhoxID and prepared three biological replicates each for the irradiated samples and nonirradiated controls. After filtering out proteins that were identified by just one peptide, we defined “hits” as proteins whose average fold change (FC) value was ≥2.5 and whose *p*-value was <0.05. Using these criteria, labelling with reagent **1** detected the target GRID2 as an enriched protein, but only 24 hits were obtained out of the 3,681 proteins detected (**Fig. 3b**, **Supplementary Data 2**). By contrast, labelling with reagents **2** and **3** enriched 164 and 243 proteins, respectively, including GRID2 and many benchmark proteins known to interact with GRID2 both intracellularly and extracellularly. Focusing on the proteins identified using **2**, for instance, CBLN1 and NRXNs are known to form a transsynaptic ternary complex with GRID2^26^. On the postsynaptic side, various intracellular scaffolding proteins such as disks large homolog 2 and 4 (DLG2 and 4, also known as PSD93 and 95), SHANK1, SHANK2, and delphilin (GRID2IP), which are known to anchor GRID2 in the postsynaptic density, were enriched (**Fig. 3b**)^32,33^. On the presynaptic side, many intracellular active zone proteins related to the release of synaptic vesicles were identified, including synaptotagmins (SYT3, SYT9), calcium-dependent secretion activator 2 (CADPS2), ELKS/Rab6-interacting/CAST family member 1 (ERC1), liprin-alphas (PPFIA2, PPFIA3), protein piccolo (PCLO), regulating synaptic membrane exocytosis protein 1 (RIMS1), protein unc-13 homolog A (UNC13A) and peripheral plasma membrane protein (CASK)^34^. CASK serves as a link between the extracellular matrix and the actin cytoskeleton and, in particular, interacts with the cytoplasmic domain of NRXNs, which in turn binds to GRID2.

**Figure 3.**
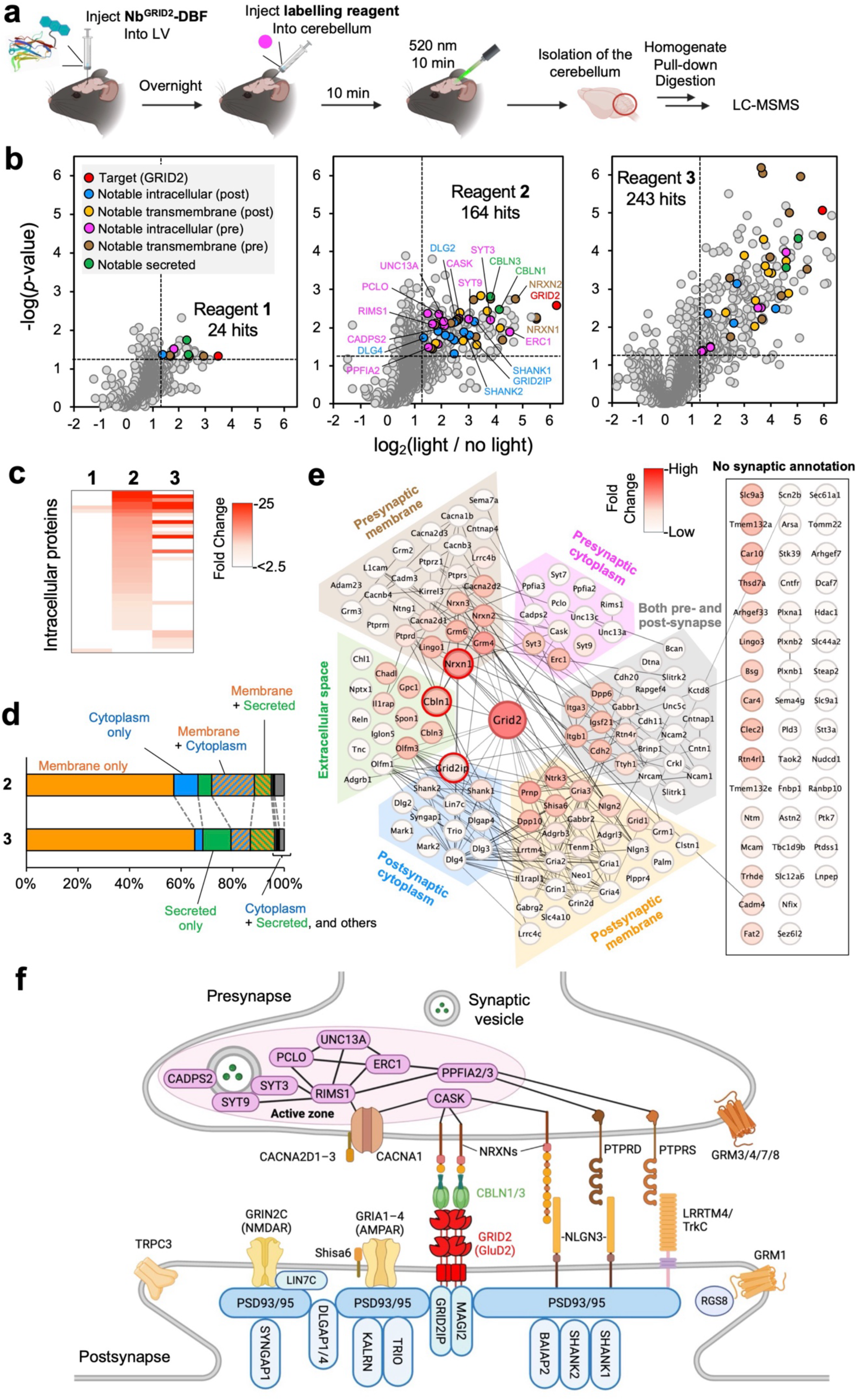
tmPhoxID labelling targeting GRID2. **(a)** Schematic overview of the experimental workflow for the *in vivo* tmPhoxID protocol. The GRID2-targeted Nb-DBF conjugate (Nb^GRID2^-DBF) is injected into both LVs of the live mouse brain. On the following day, the labelling reagent was injected into the cerebellum, and the same injection site was irradiated with an optical fibre (520 nm). The cerebellum tissue was isolated and homogenized, followed by the enrichment of labelled proteins and tryptic digestion for the LC–MSMS analysis. **(b)** Volcano plots obtained from labelling experiments using **1** (left), **2** (middle), and **3** (right). Biological replicate n = 3. **(c)** Heatmap illustrating the relative abundance of intracellular hit proteins identified with each labelling reagent. **(d)** Stacked bar chart depicting the subcellular localization of hit proteins from datasets generated with **2** and **3**, as annotated in the UniProt database. **(e)** STRING network diagram illustrating the interactions of the proteins labelled with **2**. The interactions shown are derived from experimental evidence, curated databases, and text mining. **(f)** Cartoon depiction of select proteins identified by GRID2-targeting tmPhoxID.

Although labelling with **3** resulted in a greater number of hit proteins, an analysis of intracellular protein abundance ratios indicated that tmPhoxID labelling with **2** captured a larger proportion of intracellular proteins (**Fig. 3c**). Gene Ontology (GO) enrichment analysis of the hit proteins also showed that cytoplasmic proteins (those with only cytoplasm annotations and those with both cytoplasm and membrane annotations) were more enriched upon labelling with **2**, whereas secreted proteins and membrane proteins were more prevalent in the dataset obtained with **3** (**Fig. 3d**). From these data, we concluded that **2** is best suited for tmPhoxID aimed at capturing the intracellular interactome, whereas **3** is preferable when the primary goal is labelling extracellular regions (as in the conventional PhoxID protocol)^20^. Note that our tmPhoxID is also compatible with ex vivo studies using acute brain slices, in which **1** showed excellent performance comparable to or superior to that of *in vivo* tmPhoxID with **2** (**Extended Data Fig. 4**, **Supplementary Data 3**).

To further characterize tmPhoxID with **2**, we visualized the 164 hit proteins in a STRING diagram and organized them into clusters based on their synaptic localization (**Fig. 3e**). Protein enrichment was indicated by a colour scale, with deeper red representing higher enrichment. GRID2 was shown in the most intense red, and within each cluster, the most strongly coloured proteins corresponded to the direct GRID2 interactors (NRXN1, CBLN1, GRID2IP). This result suggests that enrichment is, to some extent, correlated with proximity to GRID2 and interaction strength. Generally, TMPs and secreted proteins were identified with greater number and FC values than entirely intracellular proteins (**Fig. 3e**). This result is unsurprising because ^1^O_2_ concentrations are expected to be higher outside the cell because the photosensitizer is anchored to the extracellular domain of GRID2.

Figure 3f provides an illustration of notable GluD2-proximal proteins identified as hits in this study, showing their synaptic localization, diverse topologies, and physical interactions. Notably, the hit proteins include AMPAR and NMDAR, along with their auxiliary proteins, which are scaffolded by the postsynaptic density and thus reside near GRID2^35^. These postsynaptic neurotransmitter receptors and their intracellular scaffolding proteins have been proposed to cluster and align with presynaptic vesicles and their associated proteins, forming a transsynaptic nanocolumn to enhance neurotransmission efficiency^36^. Our GRID2-tmPhoxID dataset includes not only postsynaptic density-associated proteins but also representative proteins localized to the active zone, offering the potential to map the molecular constituents of the nanocolumn at parallel fibre–Purkinje cell synapses.

### Resolving synapse-specific microenvironments with tmPhoxID

To demonstrate the generality and selectivity of *in vivo* tmPhoxID, we next targeted GABA_A_R, the principal mediator of inhibitory neurotransmission in the brain, whose surrounding protein environments differ substantially from those of GRID2^5^. Notably, because both the N- and C-termini of the α, β, and γ subunits that constitute GABA_A_R are located on the extracellular side, GABA_A_R represents one of the challenging targets for intracellular interactome mapping using enzyme-mediated PL. To chemically anchor a photosensitizer to GABA_A_R, a ligand-directed covalent probe **7** was bilaterally injected into the LVs of live mice (**Extended Data** Fig. 5a)^37^. After confirming the successful anchoring of MBF to GABA_A_R (**Extended Data** Fig. 5b**–f**), we performed tmPhoxID with **2** in the cerebellum (**Extended Data** Fig. 6a). Out of the 4,076 detected proteins, 173 proteins were enriched with FC ≥ 1.5 and *p*-value < 0.05, including five GABA_A_R subunits (GABRG2, GABRA1, GABRA3, GABRB2, GABRB3) as well as NLGN2, GLRB, and MDGA2, which are known to localize at inhibitory synapses (Fig. 4a, **Supplementary Data 4**). Key GABA_A_R-associated proteins such as gephyrin (GPHN), vasodilator-stimulated phosphoprotein (VASP), and collybistin (ARHGEF9), were identified as intracellular interacting partners (Fig. 4b), indicating the successful implementation of GABA_A_R-tmPhoxID^38^. Notably, no intracellular presynaptic proteins were identified, presumably because of the low photosensitizer anchoring efficiency, the lower expression level of GABA_A_R compared with that of GRID2, and/or the long distance from the photosensitizer anchored to postsynaptic GABA_A_R to the presynapse.

**Figure 4.**
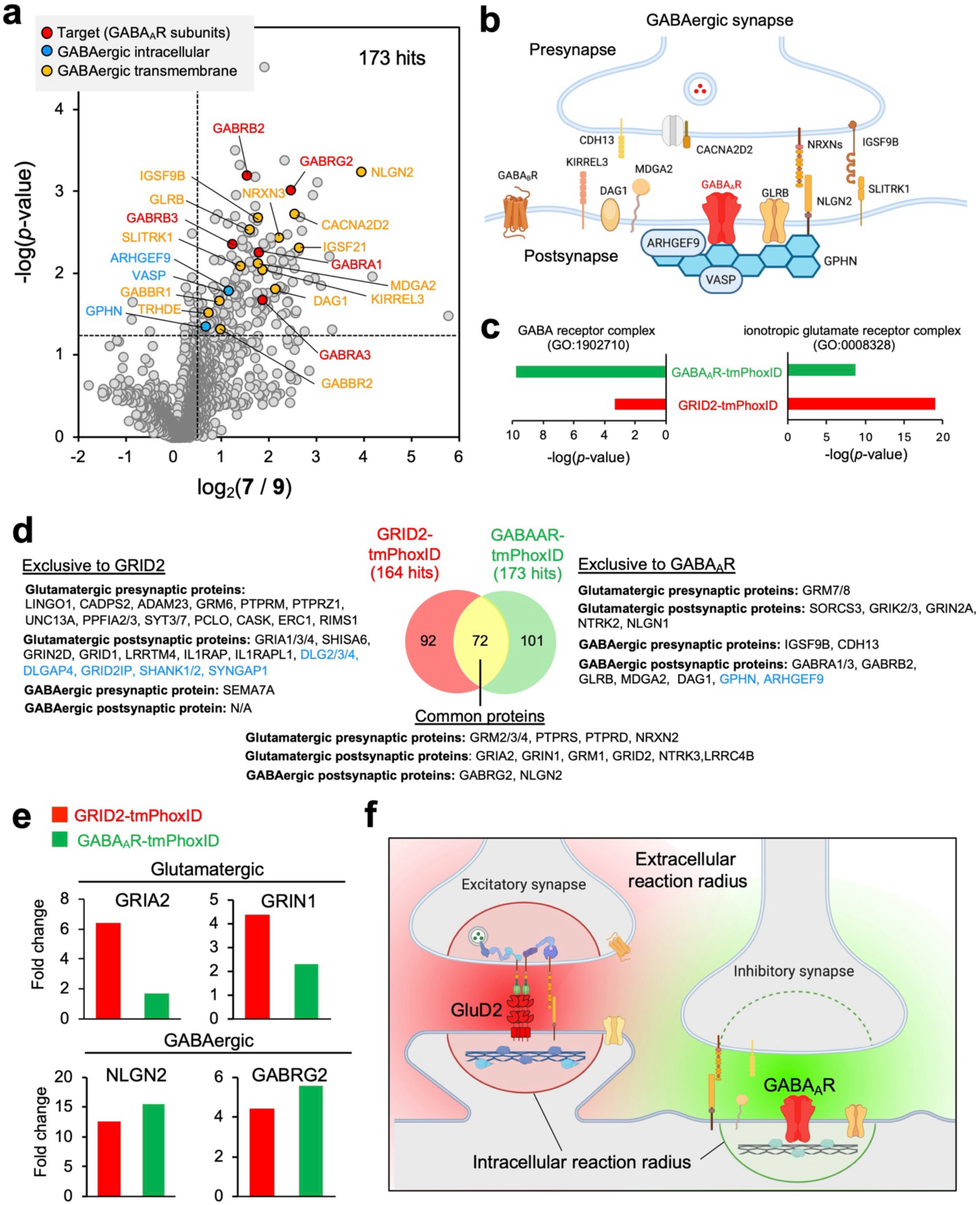
Targeting GABA_A_R to assess the spatial resolution and selectivity of tmPhoxID. **(a)** Volcano plot highlighting significantly enriched GABA_A_R-interacting proteins. Biological replicate n = 3. **(b)** Cartoon representation of the synaptic organization of GABA_A_R interactome. **(c)** GO enrichment analysis revealed that the “GABA receptor complex” term (GO:1902710) was more prominently enriched in the GABA_A_R-tmPhoxID dataset than in the GRID2-tmPhoxID dataset, while the GRID2-tmPhoxID dataset contained more enriched proteins annotated with the “ionotropic glutamate receptor complex” term (GO:008328). **(d)** Venn diagram comparing hit proteins identified in the GABA_A_R- and GRID2-tmPhoxID datasets, highlighting shared and unique proteins. **(e)** Comparison of FC values for selected glutamatergic and GABAergic proteins common to both GABA_A_R- and GRID2-tmPhoxID datasets. Glutamatergic synapse-associated proteins were more enriched in the GRID2 dataset, whereas GABAergic synapse-associated proteins showed higher enrichment in the GABA_A_R dataset. **(f)** Schematic representation of the estimated labelling radius of tmPhoxID. Red indicates the labelling radius for GRID2-tmPhoxID, with an intracellular radius confined to its pre- and postsynaptic interactome and a broader extracellular radius. Green indicates the labelling radius for GABA_A_R-tmPhoxID, with an intracellular radius restricted to postsynaptic interactors and a broader extracellular radius. Although the extracellular labelling radii of the two targets overlap to some extent, their intracellular radii remain distinct.

Comparison of the GABA_A_R- and GRID2-tmPhoxID datasets showed that the GO term “GABA receptor complex (GO:1902710)” was markedly enriched in the GABA_A_R dataset relative to GRID2, whereas the GO term “ionotropic glutamate receptor complex (GO:0008328)” was more enriched in the GRID2 dataset, showing the high target selectivity of our method (Fig. 4c). Venn diagram analysis revealed that the 92 proteins unique to the GRID2 dataset included intracellular proteins characteristic of excitatory postsynapses (DLG2/3/4, DLGAP4, GRID2IP, SHANK1/2, and SYNGAP1), whereas the GABA_A_R dataset exclusively contained intracellular proteins characteristic of inhibitory postsynapses (GPHN and ARHGEF9) (Fig. 4d). We also noted an overlap of 72 proteins between the two datasets, which included cell-surface proteins associated with both excitatory and inhibitory synapses. Notably, among these overlapping proteins, excitatory synapse–associated proteins exhibited higher FC values in the GRID2 dataset, whereas inhibitory synapse–associated proteins were preferentially enriched in the GABA_A_R dataset (Fig. 4e). Taken together, these findings characterize the reaction range of tmPhoxID as follows (Fig. 4f): for intracellular proteins, PL proceeds with higher spatial resolution (sufficient to separate synapse types) because of the confined diffusion distance of ^1^O_2_ caused by the lipid bilayer barrier and the densely crowded intracellular environment. In contrast, in the extracellular space, where the molecular density is relatively sparse, ^1^O_2_ can diffuse farther, resulting in a larger reaction radius than that inside the cell. Nevertheless, since the labelling degree strongly depends on the distance from the photosensitizer, quantitative comparison of FC enables estimation of the relative spatial relationships between specific proteins and the two target TMPs. Moreover, as demonstrated in our previous reports, extracellular PhoxID with higher spatial resolution can be achieved by adjusting the intensity and duration of light irradiation^20^.

### Resolving subsynaptic microdomains with tmPhoxID labelling

Encouraged by the high target-selectivity of tmPhoxID, we further examined whether this approach could discriminate neighbouring proteins of targets localized within the same excitatory synapse but residing in distinct subsynaptic microenvironments. To this end, we targeted the metabotropic glutamate receptor GRM1 (class C GPCR), which is known to be enriched in the perisynaptic region of excitatory synapse, and compared its interactome with that of GRID2 residing at the postsynaptic density (Fig. 5a)^39^. GRM1-targeted tmPhoxID with probe **8** yielded a proteomic profile in which GRM1 emerged as one of the most strongly enriched proteins (**Extended Data Fig. 7 and Supplementary Data 5**). Among the identified proteins, various known interactors of GRM1 were identified, including GABABR1, KCTD12, and several G proteins (GNAQ, GNAO1)^4^. Additionally, the presence of multiple extracellular and intracellular proteins associated with glutamatergic synapses further supports the successful labelling of the GRM1 interactome (**Extended Data Fig. 8**).

**Figure 5.**
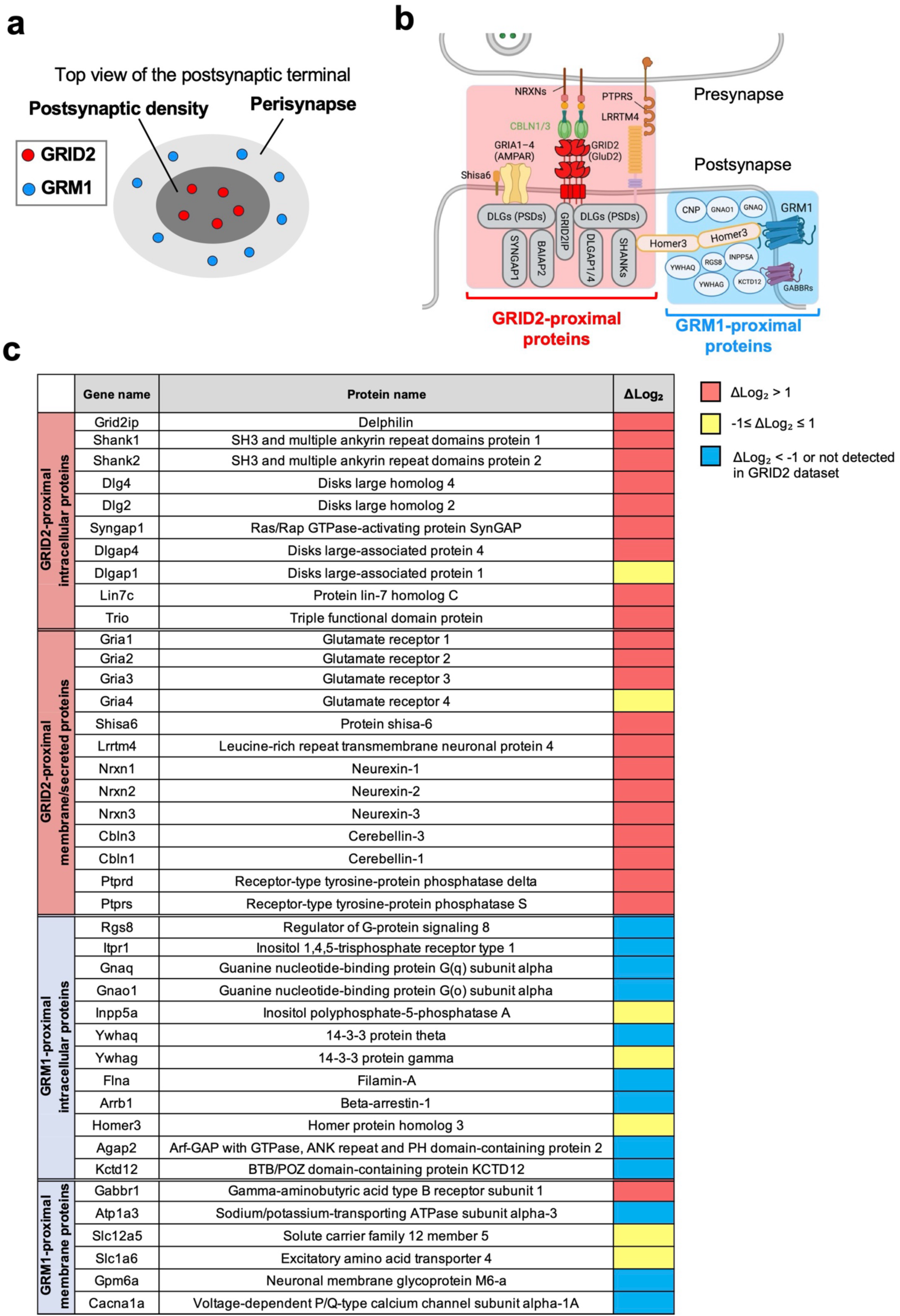
Targeting GRM1 to assess the subsynaptic spatial resolution of tmPhoxID. **(a)** Schematic representation of the targeted subsynaptic microenvironments, the perisynaptic and synaptic regions. **(b)** Schematic depiction of protein organization within the excitatory synapse, highlighting the GRM1-mediated downstream signalling cascade**. (c)** A colour map showing the tmPhoxID FC value for known neighbouring proteins of GRID2 and GRM1. Δlog2 is defined as log2 (GRID2-tmPhoxID FC) − log2 (GRM1-tmPhoxID FC).

To quantitatively assess the selectivity of GRID2- and GRM1-tmPhoxID, we compiled lists of representative known interactors for each target, as well as proteins expected to be in proximity on the basis of their synaptic localization (both extracellular and intracellular), and calculated the difference in log2 (FC) between GRID2- and GRM1-tmPhoxID (Δlog2) (Fig. 5b, c). This analysis clearly demonstrates that tmPhoxID can distinguish the unique proximal proteins of GRID2 and GRM1. For example, scaffolding proteins localized to the postsynaptic density, including GRID2IP, SHANK1/2, DLG2/4, and BAIAP2, were markedly enriched in the GRID2 dataset (Fig. 5c). By contrast, components of GRM1-mediated intracellular signalling pathways, such as G-proteins (GNAQ, GNAO1), regulator of G-protein signalling 8 (RGS8), and beta-arrestin1 (ARRB1), were present in the GRM1 dataset but absent from the GRID2 dataset^4,33^. Some proteins exhibited similar levels of enrichment in both datasets (-1 ≤ Δlog2 ≤ 1), suggesting that these proteins are proximal to both GRID2 and GRM1. Notably, the scaffolding protein HOMER3 has been reported to interact not only with GRM1 but also with SHANK1, which is near GRID2, consistent with our present observations^4,40^.

Collectively, these results indicate that tmPhoxID enables the receptor-specific labelling of proteins residing within the same synapse yet segregated into distinct subsynaptic microenvironments, highlighting the accuracy and precision of tmPhoxID labelling.

### Identification of CAMKV as a novel interaction partner of GABA_A_R

Finally, we examined whether tmPhoxID could identify previously unrecognized protein–protein interactions *in vivo*. Here, we aimed to identify intracellular proteins that interact with GABA_A_R. Candidate GABA_A_R-interacting proteins were selected on the basis of the following criteria: (1) detection with high confidence (FC > 2, *p*-value < 0.05, unique peptides ≥ 2); (2) detection exclusively in the GABA_A_R dataset but not in the GRID2 or GRM1 datasets; (3) lack of transmembrane domains; (4) no previously reported interaction with GABA_A_R; and (5) availability of antibodies for validation.

Among the four proteins that satisfied these criteria (**Supplementary Fig. 2**), we focused on CAMKV (calmodulin kinase-like vesicle-associated protein) on the basis of its high detection confidence (FC = 4.4 in cerebellum) (**Supplementary Data 4**). CAMKV is a brain-enriched pseudokinase belonging to the CaMK family^41^. Although the molecular mechanism remains unclear, recent studies suggest that CAMKV is essential in maintaining spine stability and synaptic plasticity^42,43^. Because CAMKV is more abundantly expressed in the hippocampus and cerebral cortex than in the cerebellum, we performed GABA_A_R-tmPhoxID in the hippocampus and confirmed that CAMKV was detected as a high FC (FC = 2.1 in hippocampus, **Extended Data Fig. 6 and Supplementary Data 4**).

To examine the potential interaction between CAMKV and GABA_A_R, we co-expressed CAMKV-FLAG (Fig. 6a) and the GABA_A_R subunits (α1, β3, and γ2) in HEK293T cells, followed by immunostaining. Consistent with previous reports, CAMKV localized to the plasma membrane because of its palmitoylation at Cysteine 5 (C5) and thus colocalized with GABA_A_R in the transfected cells (**Extended Data Fig. 9**)^43^. Co-IP experiments using the anti-FLAG antibody successfully pulled down the GABA_A_R subunits, strongly supporting direct interaction between CAMKV and GABA_A_R (Fig. 6b). Upon the introduction of a C5S mutation into CAMKV, which abolishes palmitoylation at this site, CAMKV showed a modest reduction in plasma membrane localization but retained detectable membrane association, and Co-IP with GABA_A_R was still observed (Fig. 6b, **Extended Data Fig. 9b**). These results indicate that the palmitoylation of CAMKV stabilizes its membrane localization but is not essential for either membrane association or interaction with GABA_A_R. To obtain more detailed structural insights into the interaction between CAMKV and GABA_A_R, we next performed deletion assays of CAMKV. Constructs lacking each of the major domains of CAMKV—the pseudokinase domain (CAMKV(ΔPK)), the calmodulin-binding domain (CAMKV(ΔCaM-BD)), or the intrinsically disordered region (CAMKV(ΔIDR))—were expressed in cells, and their interactions with GABA_A_R were evaluated by co-IP (Fig. 6a). The GABA_A_R signal was markedly reduced in CAMKV(ΔPK) and CAMKV(ΔIDR) (Fig. 6b). These findings strongly suggest that both the PK and the IDR participate in the interaction between CAMKV and GABA_A_R. Interestingly, ColabFold-based structural predictions also suggested plausible interaction interfaces between the PK of CAMKV and GABA_A_R (Fig. 6c), whereas the contribution of the IDR could not be evaluated, as expected given its lack of a defined structure. Subcellular localization analysis of these domain-deletion mutants further revealed that CAMKV(ΔPK) and CAMKV(ΔCaM-BD) exhibited completely cytosolic localization (**Extended Data Fig. 9b**). Taken together, these results indicate that the PK and the IDR of CAMKV independently contribute to its interaction with GABA_A_R, whereas the membrane localization of CAMKV requires the presence of both the PK and the CaM-BD.

**Figure 6.**
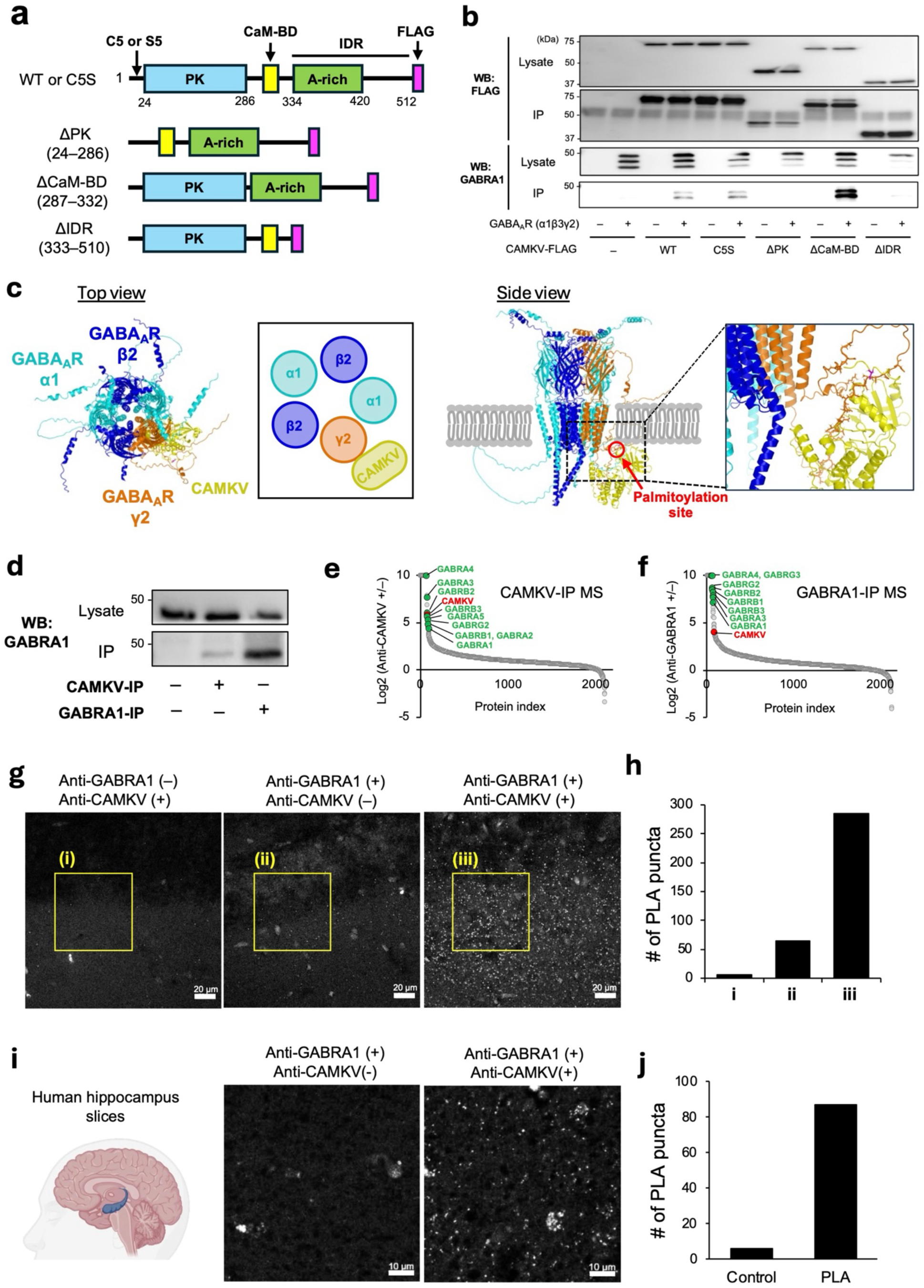
Validation of the GABA_A_R–CAMKV interaction. **(a)** Functional domain of CAMKV-FLAG. PK, pseudokinase domain; CaM-BD, calmodulin-binding domain; A-rich, alanine-rich domain; IDR, intrinsic disordered region. **(b)** Co-IP assays using HEK293T cells co-expressing CAMKV-FLAG variants and GABA_A_R. **(c)** ColabFold-predicted structural model of the complex between the CAMKV pseudokinase domain and GABA_A_R (α1β2γ2). Notably, the palmitoylation site of CAMKV is positioned adjacent to the inner-leaflet–facing side of the transmembrane helices of GABA_A_R. **(d)** Co-IP assays using mouse cerebral cortex and hippocampus lysates with anti-CAMKV and anti-GABRA1 antibodies. **(e, f)** Co-IP MS analysis of hippocampal samples using **(e)** anti-CAMKV and **(f)** anti-GABRA1 antibodies. **(g)** PLA in mouse hippocampus slices with anti-CAMKV and anti-GABRA1 antibodies as primary antibodies. **(h) N**umber of puncta in the yellow square of panel **g**. **(i)** PLA in the postmortem human hippocampus slices with anti-CAMKV and anti-GABRA1 antibodies as primary antibodies. **(j)** Quantification of number of puncta in panel **i**.

Pull-down assays using mouse hippocampus lysates with anti-CAMKV antibodies also confirmed the Co-IP of the GABA_A_R α1 subunit (Fig. 6d; the specific binding of these antibodies to their target antigens was verified in cell-based assays showing in **Extended Data Fig. 9a and Supplementary Fig. 3**). Furthermore, Co-IP MS with the anti-CAMKV antibody demonstrated enrichment of the major GABA_A_R subunits (α1–5, β1–3, and γ2) in the hippocampus neuron, whereas the pull-down assay with anti-GABRA1 antibody enriched CAMKV (Fig. 6e, f**, Supplementary Data 6**). To verify the *in situ* proximity between CAMKV and GABA_A_R, we performed a proximity ligation assay (PLA) in mouse brain slices with anti-CAMKV and anti-GABA_A_R α1 subunit antibodies^44^. Compared with the controls in which either antibody was not applied, a strong PLA signal was detected in the samples treated with both antibodies, clearly indicating the proximity (less than 40 nm) of CAMKV and GABA_A_R in the mouse brain (Fig. 6g, h, **Extended Data Fig.10**). Notably, this CAMKV–GABA_A_R proximity was also observed in human hippocampal slices (Fig. 6i, j), suggesting a conserved mechanism of neural function regulation based on this interaction in humans.

Collectively, these results highlight the utility of tmPhoxID as a powerful tool for discovering previously unrecognized protein–protein interactions *in vivo*.

## Discussion

Here, we introduced the concept of transmembrane PL, in which a catalyst anchored to one side of the plasma membrane catalyses the labelling of proximal proteins on the other side. This labelling mechanism is possible with PhoxID because, unlike the reactive species employed in other PL methods, ^1^O_2_ is cell permeable. Moreover, in PhoxID, the promiscuous oxidation of the proteins by ^1^O_2_ and the labelling reaction by nucleophilic reagents are decoupled, offering greater flexibility in terms of the design of the labelling reagent (e.g., membrane permeability). Compared with conventional PL methods, tmPhoxID has the advantage that it does not require the introduction of catalysts or enzymes into the intracellular domain of a target, which allows PL to be performed while the intracellular environment is maintained fully intact to minimize artifacts. Moreover, as demonstrated in GABA_A_R application, this approach enabled PL of the intracellular regions of even-numbered transmembrane proteins whose N- and C-termini are both exposed on the extracellular side, a scenario that is challenging to address using conventional PL strategies. The flexibility in the photosensitizer targeting strategy is another key feature of our approach. Noncovalent interactions using small-molecule ligands or antibodies (nanobodies) can be used to anchor the photosensitizer to endogenous targets. When short *in vivo* residence times or dissociation from the target pose challenges, covalent strategies—such as ligand-directed chemistry or HaloTag systems—can be utilized^37,45,46^.

GRID2-tmPhoxID successfully identified many of the constituent proteins of transsynaptic nanocolumns in genetically unedited animals^36^. Capturing such three-dimensional, transcellular protein assemblies in their native states represents a novel PL concept uniquely enabled by tmPhoxID, which offers advantages over classical synaptosome fractionation commonly used for synaptic protein characterization, including reduced contamination, minimal perturbation to protein complexes, and clear distinction between different synapse types. Among the identified hits, some are likely to be previously unannotated components of transsynaptic nanocolumns, and thus the future validation of these candidates may reveal key molecules involved in the mechanisms of efficient synaptic transmission.

The power of tmPhoxID was clearly demonstrated by the identification of the previously unrecognized CAMKV–GABA_A_R interaction *in vivo*. Although the importance of CAMKV in AMPA (α-amino-3-hydroxy-5-methyl-4-isoxazole propionic acid) receptor-mediated spine maintenance has been recognized^42^, current knowledge regarding this protein and its potential role in inhibitory synaptic transmission remains very limited. This is partly because CAMKV lacks intrinsic enzymatic activity and is largely composed of intrinsically disordered regions, which complicates its structural and functional characterization. Recent studies have indicated that CAMKV in the mouse hippocampus undergoes palmitoylation in response to context-dependent fear conditioning, leading to its reversible localization at the plasma membrane, and it interacts directly with Arc (activity-regulated cytoskeleton-associated protein) to regulate cytoskeletal organization^43,47^. Furthermore, Arc has been reported to act as a downstream effector of the small GTPase Rac1, translocating to postsynaptic regions to promote GABA_A_R endocytosis and thereby contributing to the extinction of aversive memories^48^. Combining these previous findings with our identification of the CAMKV–GABA_A_R interaction suggests that CAMKV may function as a molecular mediator linking Arc to GABA_A_Rs during Rac1-dependent GABA_A_R endocytosis. Moreover, our PLA data showed that CAMKV interacts with GABA_A_R in the human hippocampus. Interestingly, the human CAMKV gene is located on chromosome 3 (3p21.31), and copy number variations or microdeletions in this region have been reported in individuals with autism, intellectual disability, and developmental delay^42,49,50^. Thus, our findings suggest that CAMKV dysregulation may be associated with human neurological dysfunctions that arise from abnormalities in excitatory/inhibitory synapse formation, underscoring that further functional characterization of CAMKV-GABA_A_R interaction could facilitate the development of novel therapeutic strategies for neurological disorders.

A potential caveat regarding the tmPhoxID approach is the relatively large labelling radius in the extracellular region. This arises because, to deliver sufficient ^1^O_2_ into the intracellular space, tmPhoxID requires stronger light irradiation intensity than the conventional PhoxID protocol targeting only the extracellular domain. In this study, we demonstrated that the FC value can be used as an indicator for proximity to distinguish between proximal and distal proteins; however, it should be noted that inappropriate light irradiation conditions could compromise the accuracy of data interpretation.

Future tmPhoxID studies targeting other cell-surface TMPs may help characterize protein complexes spanning cell-to-cell contact sites not limited to neuronal synapses. TMPs in intracellular organelle membranes also represent important future targets, and their study could enable the comprehensive identification of three-dimensional protein complexes and networks at organelle contact sites. In addition, *in vivo* tmPhoxID is expected to be useful for tracking TMP interactomes that change over time in response to external stimuli (e.g., drug treatment) or neuronal activity, and it holds the potential to enable the simultaneous correlation of behaviours and phenotypes of individual animals with interactome profiling.

## Acknowledgments

The authors would like to thank K. Uchida, Y. Yabuki and E. Kusaka for their support in chemical synthesis and compound characterization; H. Nakamura, M. Tsujikawa and K. Matsuba for plasmid preparation; K. Nishizawa and T. Gonda for mouse experiments; and H. Utsunomiya for early-stage evaluation of nanobody-photosensitizer conjugates. We also thank Radu Aricescu and Michisuke Yuzaki for their contributions to the preparation of nanobodies. We thank Laura Murray, PhD, from Edanz for editing a draft of this manuscript. This work was supported by the Japan Society for the Promotion of Science Grant-in-Aid for Specially Promoted Research 23H05405 to I.H.; the JST FOREST program (JPMJFR230G) to T.T.; the Japan Society for the Promotion of Science Grant-in-Aid for JSPS Fellows 21J23228 to M.T.; the Ministry of Education, Culture, Sports, Science and Technology (MEXT) scholarship to F.Y.T.V.

## Author contributions

F.Y.T.V., M.T., T.T., and I.H. conceived the project and designed the experiments. F.Y.T.V., M.T., A.A., S.S., and T.T. performed the experiments and data analysis. F.Y.T.V., M.T., T.T., and I.H. wrote the manuscript with input from all authors.

## Data availability

The MS raw data and analysis files have been deposited in the ProteomeXchange Consortium (http://proteomecentral.proteomexchange.org) via the jPOST partner repository (http://jpostdb.org)^51^ with data set identifier JPST004324. All data supporting the findings of this study are available within the paper and the Supplementary Information files.

## Methods

### General materials and methods for biochemical/biological experiments

Unless otherwise noted, all proteins/enzymes and reagents were obtained from commercial suppliers (Sigma-Aldrich, Merck, Tokyo Chemical Industry (TCI), Fujifilm Wako Pure Chemical Corporation, Bio-Rad, or Thermo Fisher Scientific) and used without further purification. SDS-PAGE and western blotting were carried out using a Bio-Rad Mini-PROTEAN III electrophoresis apparatus. Samples were applied to SDS-PAGE and electrotransferred onto polyvinylidene fluoride membranes (Bio-Rad), followed by blocking with 5% nonfat dry milk in Tris-buffered saline containing 0.05% Tween 20 (TBS-T). Primary antibodies are indicated in each experimental procedure, and anti-rabbit IgG-HRP conjugate (Cell Signaling Technology, 7074S, 1:4000) or anti-mouse IgG-HRP conjugate (Cell Signaling Technology, 7076S, 1:4000) were used as the secondary antibodies. Chemiluminescent signals generated with ECL Prime (Cytiva) were detected with a Fusion Solo S imaging system (Vilber Lourmat).

### General information for fluorescence imaging experiments

Confocal laser scanning microscopy (CLSM) was performed using TCS SP8 (Leica Microsystems, Germany) or LSM 800 (Carl Zeiss AG, Germany) equipped with a 5× objective (NA = 0.15 dry objective for TCS SP8), 5× objective (NA = 0.25 dry objective for LSM 800), 10× objective (NA = 0.40 dry objective), 63× objective (NA = 1.40 oil objective), and a GaAsP detector. The excitation laser was derived from a 488 nm, 561 nm, 640 nm diode laser (LSM 800) or a white laser (TCS SP8) and was set to an appropriate wavelength depending on the dye. Airyscan mode (Carl Zeiss AG, Germany) was used for obtaining high-resolution images. Line profiles and puncta quantification were analyzed by ImageJ2.

### Animals

5-week-old C57/BL 6N (SLC) male mice under specific pathogen-free conditions were purchased from Japan SLC, Inc (Shizuoka, Japan). The animals were housed in a controlled environment (23 ± 1 °C, 12 h light/dark cycle) and had free access to food and water, according to the regulations of the Guidance for Proper Conduct of Animal Experiments by the Ministry of Education, Culture, Sports, Science, and Technology of Japan. All experimental procedures were performed in accordance with the National Institute of Health Guide for the Care and Use of Laboratory Animals, and were approved by the Institutional Animal Use Committees of Kyoto University.

### Plasmid construction

pLyn-Halo and pGPI-Halo: A plasmid encoding Lyn-EYFP and GPI-EYFP, kindly gifted from Dr. H. Nakamura (Kyoto University, Japan), was digested by AgeI and BsrGI followed by dephosphorylation of the 5’ terminus with calf intestine alkaline phosphatase (CIP). The cDNA encoded HaloTag sequence was subcloned between the AgeI and BsrGI sites of these plasmids to generate pLyn-Halo and pGPI-Halo.

pHalo-EGFR-FRB: A plasmid encoding HA-H6K-EGFR^52^ was digested by SacII and KpnI followed by dephosphorylation of the 5’ terminus with CIP. The cDNA encoded HaloTag sequence was subcloned between the SacII and KpnI sites of the plasmid to generate pHA-Halo-EGFR. The C-terminal region of EGFR and a portion of the downstream vector sequence in pHA-KH6-EGFR were excised with ClaI and XhoI and subcloned into pBluescript (pBS) to generate pBS(EGFR-C). Using pBS(EGFR-C) as a template, the stop codon at the EGFR C-terminus was removed and BsrGI/XhoI sites were introduced by PCR to obtain pBS(EGFRC-BsrGI-XhoI). Subsequently, the FRB fragment was amplified from pEGFP-FRB (kindly provided by Dr. H. Nakamura) using primers that added a BsrGI site, a linker (SAGG) at the 5′ end, a stop codon, and an XhoI site at the 3′ end. The amplified FRB fragment was then ligated into the BsrGI/XhoI-digested pBS(EGFR-C-BsrGI-XhoI) to generate pBS(EGFR-C-FRB). Finally, pBS(EGFR-C-FRB) was digested with ClaI/XhoI to excise a 1.1 kb fragment, which was ligated into the ClaI/XhoI-digested 9.1 kb fragment of pHA-Halo-EGFR to construct pHA-Halo-EGFR-FRB.

pCAMKV-FLAG: The cDNA encoding full-length mouse CAMKV-FLAG tag with NheI and NotI sites was purchased from AZENTA life sciences (pUC-CAMKV-FLAG). The cDNA was subcloned between the NheI and NotI sites of pCAGGS vector to generate pCAMKV-FLAG.

pCAMKV (C5S)-FLAG: Using pUC-CAMKV-FLAG as a template, the C5S mutation was introduced by conventional PCR with Q5 Site Directed Mutagenesis Kit (New England Biolabs) and following primers: 5’-GCCCTTCGGCtctGTGACCCTGG-3’ and 5’-ATGGTGGCGCTAGCATCC-3’.

Introduction of the C5S mutation was verified by DNA sequencing. The cDNA fragment digested by NheI and NotI was inserted into pCAGGS vector to generate pCAMKV (C5S)-FLAG. pCAMKV(ΔPK)-FLAG, pCAMKV(ΔCaM-BD)-FLAG, pCAMKV(ΔIDR)-FLAG: Using pUC-CAMKV-FLAG as a template, the peudokinase domain (24–286), the calmodulin binding domain (287–332), the intrinsic disordered region (333–510) were deleted by conventional PCR with Q5 Site Directed Mutagenesis Kit and following primers: 5’-AGCGGCAACGCCGCCTCC -3’ and 5’-TCTGTCGGTCACCTCGCTAGGC -3’ for ΔPK; 5’-GGGACAGCTGCCACACAG-3’ and 5’-GATCCACTCGTGGCTGATG -3’ for ΔCaM-BD; 5’-ACAAGCGGCGGGGGCAGC-3’ and 5’-GCTTTGCTCGGGAGCTCTCAGTCTC -3’ for ΔIDR. Deletion was verified by DNA sequencing. The cDNA fragments digested by NheI and NotI were inserted into pCAGGS vector to generate pCAMKV(ΔPK)-FLAG, pCAMKV(ΔCaM-BD)-FLAG, and pCAMKV(ΔIDR)-FLAG.

### Cell culture and transfection

HEK293T cells (purchased from ATCC) were cultured in high glucose Dulbecco’s Modified Eagle Medium (DMEM, 4.5 g of glucose/L) supplemented with 10% fetal bovine serum (FBS), 1X Antibiotic-Antimycotic (gibco). Cells were maintained at 37°C in a humidified atmosphere of 5% CO_2_ in air. A subculture was performed every 3-4 days from subconfluent (<80%) cultures using a trypsin-EDTA solution. Lipofectamine 2000 (Thermo Fisher Scientific) was used for plasmid transfection according to manufacturer’s instructions. Cells 18–24 h post-transfection were used for experiments.

### Labelling experiment with Lyn-Halo

HEK293T cells transfected with pLyn-Halo were treated with **AcDBF-HL** (**4**) (1 µM) in DMEM-HEPES (FBS(–)) for 30 min at 37 °C in a 5% CO_2_ incubator. The cells were washed twice with DMEM-HEPES (FBS(–)), 30 min each at 37 °C in a 5% CO_2_ incubator, then treated with **Hyd-PEG4-Bt** (**6**) (1 mM) in HEPES-buffered saline (HBS) buffer (20 mM HEPES, 107 mM NaCl, 6 mM KCl, 2 mM CaCl_2_, 1.2 mM MgSO_4_, 11.5 mM Glucose, pH7.4) for 10 min at r.t. After irradiation with a homemade LED irradiation device (lambda max 510 nm, irradiance= 11–19 W/m^2^, described previously^21^) for 5 min, cells were kept in darkness for 5 min, washed thrice with phosphate-buffered saline (PBS(–)). Streptavidin-AF647 (Invitrogen, 2 µg/mL) in PBS(–) was added to the dishes for 5 min at room temperature, then washed with PBS(–) twice before imaging with the LSM 800.

### Rapamycin-induced labelling of mCherry-FKBP12

HEK293T cells co-transfected with pHaloTag-HA-EGFR-FRB and pmCherry-FKBP12 were treated with **MBF-HL** (**5**) (1 µM) in DMEM-HEPES (FBS (–)) for 30 min at 37°C in a 5% CO_2_ incubator. The cells were washed twice with DMEM-HEPES (FBS (–)), 30 min each at 37°C in a 5% CO_2_ incubator, then treated with **1**, **2**, or **3** (1 mM) in HBS for 20 min at r.t. in the presence or absence of rapamycin (5 µM). Cells were then irradiated for 5 min using a homemade LED irradiation device on top of aluminum foil with the dish lids removed or kept in darkness for the same duration. After 5 min in darkness, the samples were collected into 1.5-mL tubes and washed thrice with PBS (–), centrifuging at 1000 g for 3 min each time. The resultant pellet was then lysed with RIPA buffer (25 mM Tris–HCl, 150 mM NaCl, 1% Nonidet P-40, 0.1% sodium dodecyl sulfate, 0.25% sodium deoxycholate, pH 7.4) containing 1% protease inhibitor cocktail set III (Millipore, 539134). The labelled proteins in the lysate were then pulled-down with High Capacity NeutrAvidin Agarose beads (Thermo Fisher Scientific), as described below. The input and the bead eluate were analyzed by western blot with anti-mCherry antibody (abcam, rabbit, ab167453, 1:2000).

### Labelling experiment with GPI-Halo

HEK293T cells transfected with pGPI-Halo were treated with **MBF-HL (5)** (1 µM) in DMEM-HEPES (FBS (–)) for 30 min at the at 37 °C in a 5% CO_2_ incubator. The cells were washed twice with DMEM-HEPES (FBS (–)), 30 min each at 37°C in a 5% CO_2_ incubator, then treated with **1** (1 mM) in HBS for 10 min at r.t. After irradiation with a homemade LED irradiation device for 5 min, cells were kept in darrkness for 5 min. After washing thrice with PBS (–), the cells were fixed and permeabilized with 4% PFA/0.1% Triton X-100/PBS (–)) for 30 min at r.t.. The cells were subsequently washed thrice with PBS (–) (7 min each), blocked with bivine serum albumin (BSA, 1 mg/mL in TBS-T) for 30 min at r.t., then washed again with PBS (–) (5 min each). The cells were then treated with a solution of 1 mM CuSO_4_, 2 mM tris(3-hydroxypropyltriazolylmethyl)amine (THPTA), 10 mM sodium ascorbate, and 15 µM Alexa Fluor 647-azide (Thermo Fisher Scientific) for 1 h at 37 °C in the dark. The reagents were then washed thrice with TBS-T, thrice with PBS (–), and kept overnight in PBS (–) for 4 °C and imaged the following day.

### Anchoring of photosensitizers in the mouse brain

C57BL/6N mice (male, 5 weeks old, body weight = 18–23 g) were anesthetized with a mixture of Domitor® (Nippon Zenyaku Kogyo Co., Ltd.), midazolam (Sandoz), and Vetorphale® (Meiji Seika Pharma Co., Ltd.) by intraperitoneal injection, and a solution of the **Nb^GRID2^-DBF^20^** (4.4 µL, 40 µM, PBS(–)), **7**, **8** or **9** (4.4 µL, 100 µM, PBS (–), 5% DMSO) was directly injected into the left and right LVs (AP = -1.3 mm from bregma, ML = ± 2 mm, depth = 2 mm) using a microinjector (Nanoliter 2010, World Precision Instruments) (600 nL/min) via burr holes drilled through the skull. After 2 min, the injection capillary was withdrawn, the scalp was clipped back together, and the anesthesia antagonist Antisedan® (Nippon Zenyaku Kogyo Co. Ltd., 0.2 mL) was administered by intraperitoneal injection. Mice were kept in their cages for 20 – 24 h post-injection to allow the binding and/or covalent anchoring reaction to proceed. For western blot-based confirmation of the covalent anchoring reaction with GABA_A_R, mice were sacrificed under deep anesthesia with isoflurane, and the mouse brain tissue was isolated, dissected, and lysed with 1% SDS RIPA buffer (pH 7.4, 25 mM Tris–HCl, 150 mM NaCl, 1% SDS, 1% Nonidet P-40, 0.25% deoxycholic acid) containing 1% protease inhibitor cocktail set III. Lysates were further homogenized with an ultrasonic cell disruptor (Branson Ultrasonics, Sonifier SFX 250). Following centrifugation (17730 g, 10 min, 4 °C), the protein concentrations of the supernatant were analyzed by BCA assay (Pierce), and the normalized lysates were mixed with a quarter volume of 5 × sample buffer (pH 6.8, 312.5 mM Tris–HCl, 25% sucrose, 10% SDS, 0.025% bromophenol blue) containing 250 mM dithiothreitol (DTT) and vortexed for 1 h at room temperature. SDS-PAGE and western blotting analyses were performed as described above. The covalently modified receptor (GABA_A_R) was detected by in-gel fluorescence analysis following SDS-PAGE and the detection of GABRG2 was conducted with rabbit anti-GABA-A receptor gamma2 (Synaptic Systems, 224003, 1:2000) western blotting.

### tmPhoxID in the live mouse brain

Mice injected with photosensitizer-anchoring reagents were anesthetized by intraperitoneal injection of a mixture of Domitor®, midazolam, and Vetorphale®. For photoinduced labelling in the hippocampus, burr holes were made at both sides and **2** (4.4 µL, 5 mM, PBS (–), 5% DMSO) was injected at AP = - 2.3 mm from bregma, ML = ±1.6 mm, depth = 1.4 mm using a microinjector at a rate of 600 nL/min. For photoinduced labelling in the cerebellum, a burr hole was made and **1** (4.4 µL, 100 mM, PBS (–)), **2** or **3** (4.4 µL, 5 mM, PBS (–), 5% DMSO) was injected into the vermis of cerebellar lobules V-VIII (AP = -8 mm from bregma, ML = +1 mm) at a depth of 0.75 mm from the tissue surface. Immediately after the end of injection, an optical fiber (Doric Lenses, 520 nm laser, LDFLS-520/060, fiber cannula: FOC-C-200.1.25-0.22-3.0) was inserted into brain tissue at the same depth as the injection through the burr hole. At 10 min after the end of injection, the light source was turned on for 10 min to irradiate the brain. Irradiance was 0.96 W/cm^2^ for all *in vivo* tmPhoxID experiments unless specified otherwise. 5 min after the end of irradiation, the mouse was sacrificed, and its brain was isolated and submerged in ice-cold cutting solution (120 mM choline chloride, 3 mM KCl, 8 mM MgCl_2_, 1.25 mM NaH_2_PO_4_, 28 mM NaHCO_3_, 22 mM glucose, 0.5 mM ascorbic acid). The hippocampus or the cerebellar vermis were macroscopically dissected and retrieved. Isolated tissues were homogenized with a pestle in 400 µL of 1% SDS-RIPA buffer containing 1% protease inhibitor cocktail set III. The lysates were further homogenized with an ultrasonic cell disruptor, and the lysates were diluted to a final SDS concentration of 0.5% for enrichment of the labelled proteins, as described below.

### Enrichment of labelled proteins

Following centrifugation (17730 g, 10 min, 4 °C) of the lysates, the protein concentration of the supernatant was determined by BCA assay. The solution was diluted to a protein concentration of 1.5 mg/mL by adding ∼0.5% SDS RIPA buffer, and 0.7–2 mg protein was added to High Capacity NeutrAvidin Agarose beads (bed volume: 100 µL) pre-washed with PBS (–) in a 1.5 mL Protein LoBind tube (Eppendorf). After rotating at r.t. for 2 h, the beads were centrifuged (3000 g, 1 min, 4 °C) and washed with RIPA buffer (×4). The beads were transferred to a fresh 1.5-mL Protein LoBind tube and further washed with RIPA buffer (×4). The beads were then mixed with 100 µL of RIPA buffer supplemented with 4 mM biotin and incubated at 37 °C for 1 h, then mixed with a quarter volume of 5× sample buffer containing 250 mM DTT and heated at 95°C for 5 min (for GRID2- and GRM1-targeted tmPhoxID) or at 45°C for 1h (for GABA_A_R-targeted tmPhoxID). After protein elution, the beads were removed using a micro bio-spin column (Bio-Rad), and the eluate was collected. A streptavidin-HRP conjugate (Invitrogen, S911, 1:4000) was used to detect the labelled proteins by western blotting.

### Sample preparation for NanoLC-MS/MS

Enriched protein samples were loaded to a 10% SDS-PAGE gel (Bio-Rad, Mini-PROTEAN TGX Gels) and resolved for ∼1 cm. The gel containing protein samples was manually excised, diced, and fixed with 45% methanol/water containing 5% acetic acid for 20 min. The fixed gel pieces were washed with 50% methanol aq. and pure water, then dehydrated with acetonitrile. The dehydrated gels were swelled with 200 µL of 10 mM DTT in 100 mM triethylamine bicarbonate (TEAB) buffer, pH 8.5 and heated at 56 °C for 30 min. The DTT solutions were replaced with 55 mM iodoacetamide in 100 mM TEAB buffer, and the gels were incubated at 37°C in the dark for 30 min. The gels were then dehydrated in acetonitrile, rehydrated in 100 mM TEAB buffer, and dehydrated in acetonitrile again. Next, the gel was swelled in 100 mM TEAB buffer containing 10 ng/µL Sequence Grade Trypsin (Promega, V5113). After incubation overnight at 37°C, an extraction solution (50% acetonitrile containing 0.1% trifluoroacetic acid, TFA) was added to the gels and the gels were sonicated for 10 min in a bath sonicator. The supernatant was collected in a new Protein LoBind tube, and this process was repeated once with 50% acetonitrile containing 0.1% TFA and twice with 100% acetonitrile containing 0.1% TFA. The collected peptide solution was concentrated by a centrifugal concentrator (TOMY CC-105) and the extracted peptides were further purified by GL-Tip SDB (GL Sciences) according to the manufacturer’s instructions.

### NanoLC-MS/MS analysis

NanoLC-MS/MS analyses in data-independent acquisition (DIA) mode were performed with the Orbitrap Exploris 480 mass spectrometer (Thermo Fisher Scientific) connected to a Vanquish Neo UHPLC system (Thermo Fisher Scientific). The peptides were loaded into a nano-HPLC capillary column (C18 packed with 3-µm gel particles, 0.075 × 125 mm, Nikkyo Technos, Tokyo Japan) via PepMap Neo trap cartridge (Thermo Fisher Scientific). The injection volume was 2 µL and the flow rate was 300 nL/min. The mobile phases consisted of (A) 0.5% acetic acid and (B) 0.5% acetic acid and 80% acetonitrile. A multi-step linear gradient of 5–5% B over 3.6 min, 5–19% B over 40.7 min, 19–29% B over 15.5 min, 29–45% B over 10.2 min, 45–95% B over 2 min, 95% B for 18 min was employed.

A spray voltages of 1,600 V, an ion transfer tube temperature of 275°C, and a normalized HCD collision energy of 30% were applied. The precursor mass scan ranges and the isolation window were set to m/z 500–860 and 6, respectively. The orbitrap resolution was set to 30000. The software package DIA-NN version 1.8.1 was used to analyze the raw MS data files. Precursor ions were generated with FASTA and deep-learning based spectra, and trypsin specificity that allowed for 1 missed cleavage against Mus Musculus, UniProt/SWISS-PROT (release 2023-10). Peptides lengths between 5 to 30 residues were tolerated. The precursor charge range was set to 1–4, the precursor *m*/*z* range to 500–860, and the fragment ion *m*/*z* range to 200–1500. Match between runs (MBR) was implemented and isotopologs were used. Precursor FDR was set to 1% with N-terminal methionine excision and C-terminal carbamidomethylation as fixed modification. Protein grouping was performed using a highly heuristic mode to minimize redundancy and reduce the total number of reported protein groups for benchmarking protein identification numbers and enable more consistent comparisons across datasets. Technical duplicates were measured for all samples in label-free quantification analysis. Protein abundances in each raw file were normalized across samples on the abundances of three known endogenously biotinylated proteins (UniProt IDs: Q99MR8, Q05920, Q91ZA3) and missing values were replaced with random values sampled from the lowest 5% of all detected values in each raw file. Geometric means of the technical replicates were calculated to obtain the protein abundance values for each biological sample (n = 3). The AlphaPeptStats Python package was used to remove common mass spectrometry contaminants and perform statistical analyses^53^.

### Photoinduced proximity labelling in acute brain slices

C57BL/6N mice (5 weeks old, male) were anesthetized with a mixture of Domitor®, midazolam, and Vetorphale®, and injected to the left and right LVs with **Nb^GRID2^-DBF** (40 µM in PBS (–), 4.4 µL on each side). Sagittal cerebellum slices (250 µm thick) were prepared with a vibratome (DOSAKA EM) and immersed in a solution of **1** (4 mM) in artificial cerebrospinal fluid (ACSF: 125 mM NaCl, 2.5 mM KCl, 2 mM CaCl_2_, 1 mM MgCl_2_, 26 mM NaHCO_3_, 1.25 mM NaH_2_PO_4_, 10 mM glucose) for 30 min at room temperature in a 35-mm glass-bottom dish (Matsunami) under 95% O_2_/5% CO_2_. The dishes containing the slices were then placed on aluminum foil and illuminated for 30 min at room temperature using a homemade photoirradiation device or kept in darkness for the same duration. After photoirradiation, the slices were washed thrice with carboxygenated ACSF (1 mL) and once with PBS (–) on ice. For each condition, 5–9 slices were pooled and lysed together in 300-400 µL of 1% SDS RIPA buffer containing 1% protease inhibitor cocktail set III. The lysate was further homogenized with an ultrasonic cell disruptor (Branson Ultrasonics, Sonifier SFX 250) and was diluted with an equal volume of 0.1% SDS RIPA buffer to reduce the SDS concentration to ∼0.5%.

### Click reaction in cell and tissue lysates for samples treated with 1

The cell or tissue lysate labelled with **1** was centrifuged (20030 g, 10 min, 4 °C) and the protein concentration of the supernatant was determined by a BCA assay. The solution was diluted to a concentration of 0.7 – 2 mg/mL with RIPA buffer. A cocktail of copper-catalyzed azide-alkyne cycloaddition (CuAAC) reagents was then added to the lysate ([CuSO_4_]_final_ = 1 mM, [biotin-PEG3-azide]_final_ = 50 µM, [tris((1-benzyl-4-triazolyl)methyl)amine]_final_ = 100 µM) and vortexed briefly before adding tris(2-carboxyethyl)phosphine (final concentration = 1 mM) and vortexing for an additional 1 h at r.t.. Samples subjected to CuAAC were treated with 4 equivalents of methanol, 1 equivalent of chloroform, and 3 equivalents of ultrapure water to precipitate the protein. The solution was centrifuged (20030 g, 5 min, 4°C) and the top layer was removed before adding an additional 4 equivalents of methanol and centrifuging (20030 g, 10 min, 4°C). After discarding the supernatant, protein precipitates were dried with N_2_-flow before proceeding to the resolubilization of the proteins. 800 µL of 0.5% SDS RIPA buffer was added to each sample and were homogenized with an ultrasonic cell disruptor and heated at 45°C for 45 min for posterior enrichment of the proteins as described above.

### Immunostaining fixed HEK293T cells

HEK293T cells were co-transfected with FLAG-tagged CAMKV and GABA_A_R (α1β3γ2) or transfected with CAMKV mutated variants (C5S, ΔPK, ΔCaM-BD, or ΔIDR). Cells were fixed with 4% paraformaldehyde (PFA) in PBS (–) for 30 min at room temperature, followed by two washes with PBS (–). Cells were then permeabilized with 0.1% Triton X-100 in PBS (–) for 15 min and blocked with 10% normal goat serum (NGS) in PBS (–) for 60 min at room temperature. After washing with PBS (–), cells were incubated with mouse anti-CAMKV (Santa Cruz, SC-517082, 1:500) or mouse anti-FLAG (Merck, F1804, 1:1000) and/or rabbit anti-GABRA1 (Sigma Aldrich, 06-686, 1:1000) primary andibodies diluted in 1% NGS/PBS (–) at 4 °C overnight. Following three washes with PBS (–), cells were incubated with Alexa Fluor 546 goat anti-mouse IgG (H+L) (Invitrogen, A11030, 1:1000) and Alexa Fluor 633 conjugate goat anti-rabbit IgG (H+L) (Invitrogen, A21070, 1:1000) secondary antibodies diluted in 1% NGS/PBS (–) at 4 °C overnight.

### Preparation of paraformaldehyde-fixed brain slices

C57BL/6N mice (male, 5 weeks old, body weight = 18–23 g) were transcardially perfused with ice-cold 4% PFA in PBS (–) (pH 7.4) (60 mL) under deep anesthesia with isoflurane. After isolation, the mouse brain was further fixed with 4% PFA/PBS (–) overnight at 4°C. After washing with PBS (–) (×3), the brain was sagittally sliced along the midline into halves and immersed in 30% sucrose/PBS (–) at 4°C over 1 – 3 days.

The brain tissues were embedded in optimal cutting temperature (OCT) compound, frozen, and sagittally sliced into 20 μm-thick sections using a cryostat (Leica Biosystems, CM1950).

### Immunostaining fixed brain slices

For GPHN immunostaining of brain slices containing labelled GABAAR (mice injected with **7**), and GABRA1 or CAMKV immunostaining of untreated brain samples, slices prepared as described above were mounted on glass slides and activated by incubation with 0.2% pepsin (pH 4) for 10 min at 37 °C in a humidified chamber. For GRM1 immunostaining, slices were not pre-treated prior to the permeabilization and blocking. Slices were permeabilized with 0.1% Triton X-100/PBS (-) for 15 minutes at room temperature and after PBS (-) washing, blocked with 2% BSA 2% NGS in PBS (–) containing 0.2% Triton X-100 for 30 min at room temperature. The primary antibody reaction was conducted with mouse anti-GPHN (SC-25311, 1:500), anti-GABRA1 (Sigma Aldrich, 06-686, 1:500), anti-CAMKV (Santa Cruz , SC-517082, 1:500) or rabbit anti-GRM1 (Frontier Institute, MSFR 104030, 1:1000) in PBS(–) containing 0.1% Triton X-100 overnight at 4 °C. The secondary antibody reaction was conducted with Alexa fluor 555 goat anti-mouse conjugate (Invitrogen, A21422, 1:200) or Alexa Fluor 546 conjugate goat anti-rabbit (Invitrogen, A11035, 1:200) in PBS (–) containing Triton X-100 for 1 h at room temperature.

### Proximity ligation assay

Protein–protein proximity was assessed using the Duolink® in situ proximity ligation assay (PLA; Sigma-Aldrich) according to the manufacturer’s instructions. PFA-fixed mouse brain slices (20 μm thick) and postmortem human hippocampal samples paraffin sections (5 µm thick, BioChain, Cat# T2234052) were mounted on glass slides and were activated by autoclaving at 120 °C in citrate buffer (pH 6) containing 0.05% Tween 20 for 20 min. After conducting PBS (-) washes (x3), slices were then encircled with a hydrophobic barrier using a PAP pen (Daido Sangyo Co., Ltd.). Slices were blocked using Duolink® blocking solution and incubated with rabbit anti-GABRA1 (Sigma Aldrich, 06-686, 1:500 in Duolink antibody diluent) and mouse anti-CAMKV (Santa Cruz, SC-517082, 1:500 in Duolink antibody diluent) at 4°C overnight. Species-specific PLA probes conjugated to oligonucleotides (PLUS and MINUS) were then applied, followed by ligation and amplification. Amplified PLA signals were detected using Texas-red labelled oligonucleotides and visualized by CLSM. Negative controls lacking one primary antibody were included to assess background signal.

### CAMKV-GABRA1 Co-IP in HEK293T cells and Co-IP MS of mouse brain samples

For CAMKV-GABA_A_R Co-IP in HEK293T cells, FLAG-tagged CAMKV variants (wild type, C5S, ΔPK, ΔCaM-BD, and ΔIDR) and GABA_A_R (α1β3γ2) were co-expressed. Cells were homogenized in 400 µL of 1% Maltose-neopentyl glycol (MNG) buffer (25 mM HEPES, 150 mM NaCl (pH 7.4) containing 1% protease inhibitor cocktail set III with an ultrasonic cell disruptor, 10 shots. Following centrifugation (17730 g, 15 min, 4 °C), the protein concentration of the supernatant was determined by BCA assay. The solution was diluted to a protein concentration of 1.5 mg/mL, and 1 mg protein was added to anti-FLAG tag antibody magnetic beads (Fujifilm, 2.5 mL, 017-25151) pre-washed with 1% MNG buffer in a 1.5 mL Protein LoBind tube. Beads were incubated with gentle rotation at 4°C for 4 hours and washed with 0.1% MNG buffer 4 times by placing the tubes on a magnetic stand. Pulled down proteins were eluted by the addition of 50 µL FLAG peptide solution (250 µg/mL in 1× TBS) supplemented with 2× Laemmli sample buffer containing 100 mM DTT, followed by heating at 95 °C for 5 min. The input and the bead eluate were analyzed by western blot with mouse anti-FLAG (Merck, F1804, 1:2000) and rabbit anti-GABRA1 (Sigma Aldrich, 06-686, 1:2000).

For the Co-IP MS of mouse brain samples, C57BL/6N mice (male, 5 weeks old) were anesthetized with isoflurane and sacrificed. Brain was isolated and submerged in ice-cold cutting solution. The hippocampus was macroscopically dissected and retrieved. Isolated tissues were homogenized with a pestle in 800 µL of 1% MNG buffer containing 1% protease inhibitor cocktail set III. The lysates were further homogenized with an ultrasonic cell disruptor, 20 shots, thrice. Following centrifugation (17730 g, 10 min, 4 °C), the protein concentration of the supernatant was determined by BCA assay. The solution was diluted to a protein concentration of 1.5 mg/mL by adding 1% MNG buffer. 1 mg of protein was incubated with Protein G Sepharose 4 Fast Flow beads (Cytiva, 17061801) pre-washed with lysis buffer (1 mL ×2) and rotated for 2 h at 4 °C. Beads were pelleted by centrifugation (3000 g, 1 min), and the supernatant was transferred to a fresh low-binding tube. Primary antibodies were added at a 1:100 dilution (rabbit anti-GABRA1, Sigma-Aldrich, 06-868; or mouse anti-CAMKV, Santa Cruz, SC-517082). Samples incubated without primary antibody served as negative controls. Lysates were rotated overnight at 4 °C and subsequently transferred to fresh tubes containing pre-washed Protein G Sepharose beads, followed by incubation for an additional 2 h at 4 °C with rotation. Beads were collected by centrifugation (3000 g, 1 min), and the supernatant was discarded. Beads were washed with lysis buffer (1 mL ×4), transferred to fresh tubes, washed an additional four times with lysis buffer, and finally pelleted by centrifugation (3000 g, 1 min). Protein elution was conducted as described for Co-IP in HEK293T cells, and the input and the bead eluate were analyzed by western blot with mouse anti-CAMKV (Santa Cruz, SC-517082, 1:2000) and rabbit anti-GABRA1 (Sigma Aldrich, 06-686, 1:2000) antibodies. Brain samples were further processed and analyzed by nanoLC-MSMS as described above.

## Extended Data Figures

**Extended Data Fig. 1.**
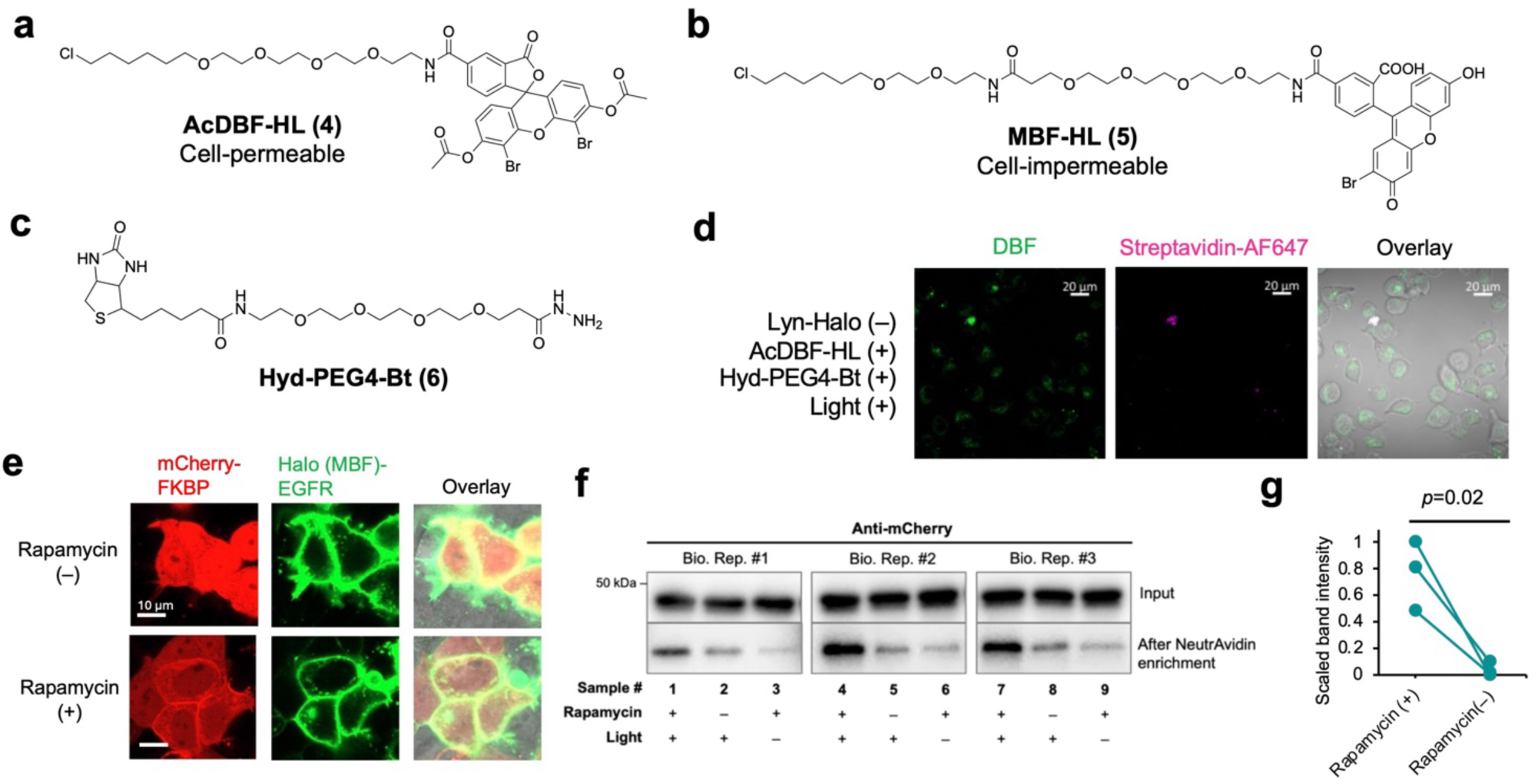
tmPhoxID in cultured cells. (**a–c**) Molecular structures of (**a**) AcDBF-HL (**4**), (**b**) MBF-HL (**5**), and (**c**) Hyd-PEG4-Bt (**6**). (**d**) Negative control for Figure 2b using cells that do not express Lyn–Halo. (**e**) Confirmation of rapamycin-dependent recruitment of mCherry–FKBP12 to the inner leaflet of the plasma membrane. (**f**) Biological replicates of pulldown detection of biotin-labelled mCherry–FKBP12. (**g**) Band intensities in the presence and absence of rapamycin. The data represent the band intensities of (**f**) obtained from each biological replicate (n = 3). Student’s one-tailed paired *t*-test was applied.

**Extended Data Fig. 2.**
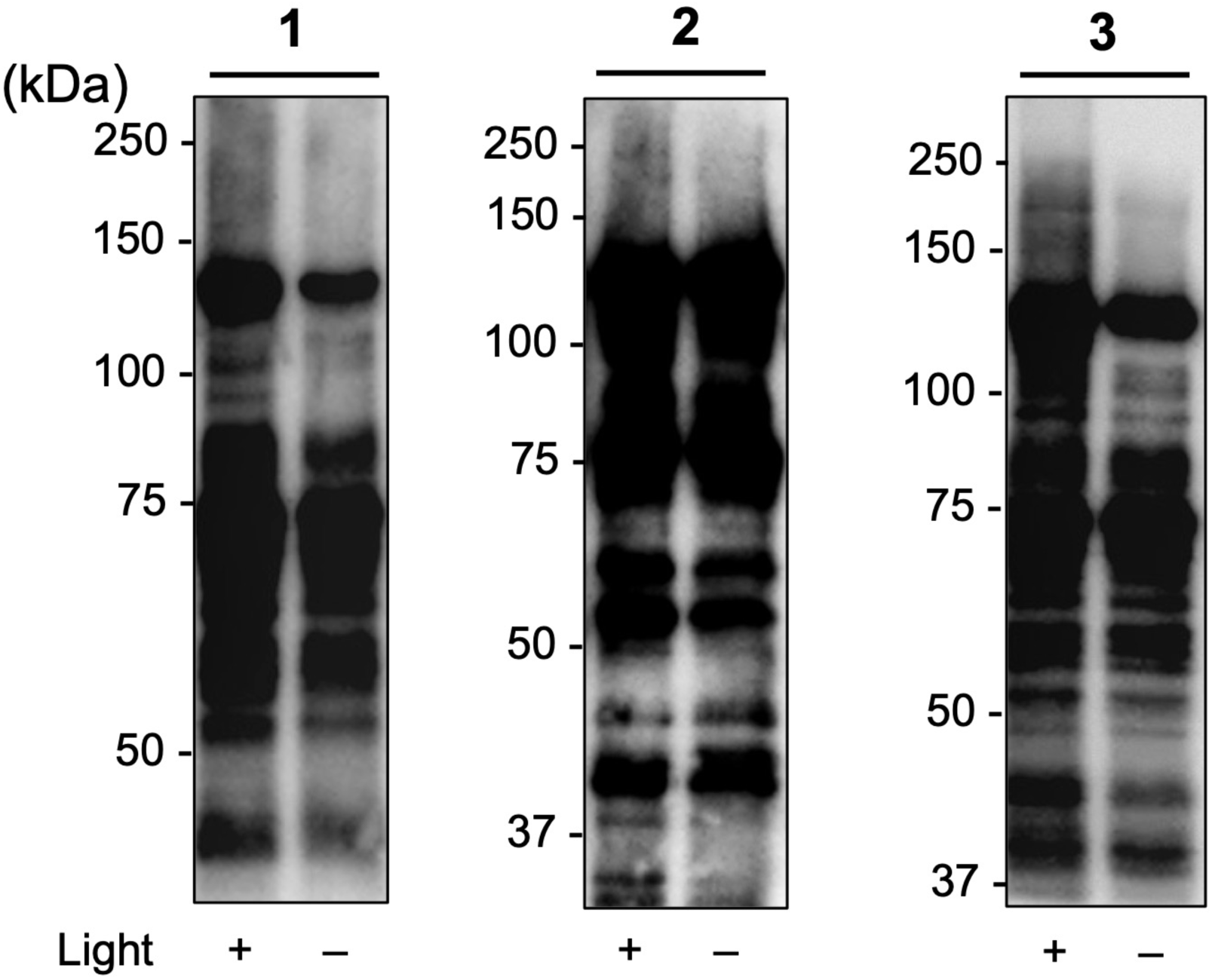
Streptavidin blot of cerebellum proteins isolated and enriched from mice subjected to *in vivo* GRID2-tmPhoxID using different labelling probes (1–3). All experiments shown are representative of at least two independent measurements.

**Extended Data Fig. 3.**
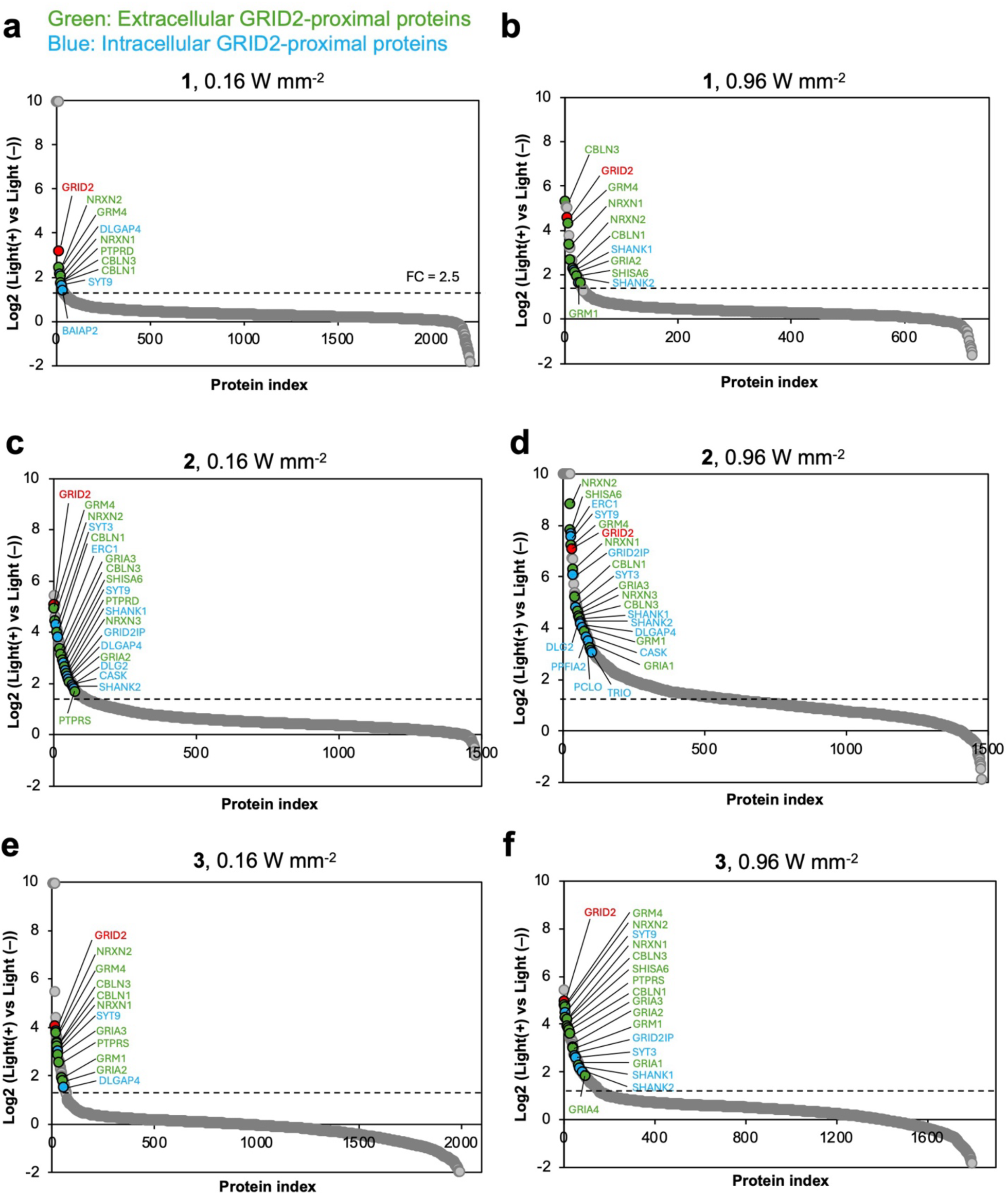
Light intensity dependence of tmPhoxID results. *In vivo* GRID2-tmPhoxID was performed with (**a**) probe **1** and irradiation at 0.16 W mm^-2^, (**b**) probe **1** and irradiation at 0.96 W mm^-2^, (**c**) probe **2** and irradiation at 0.16 W mm^-2^, (**d**) probe **2** and irradiation at 0.96 W mm^-2^, (**e**) probe **3** and irradiation at 0.16 W mm^-2^, (**f**) probe **3** and irradiation at 0.96 W mm^-2^. Light irradiation was applied for 10 min for all samples.

**Extended Data Fig. 4.**
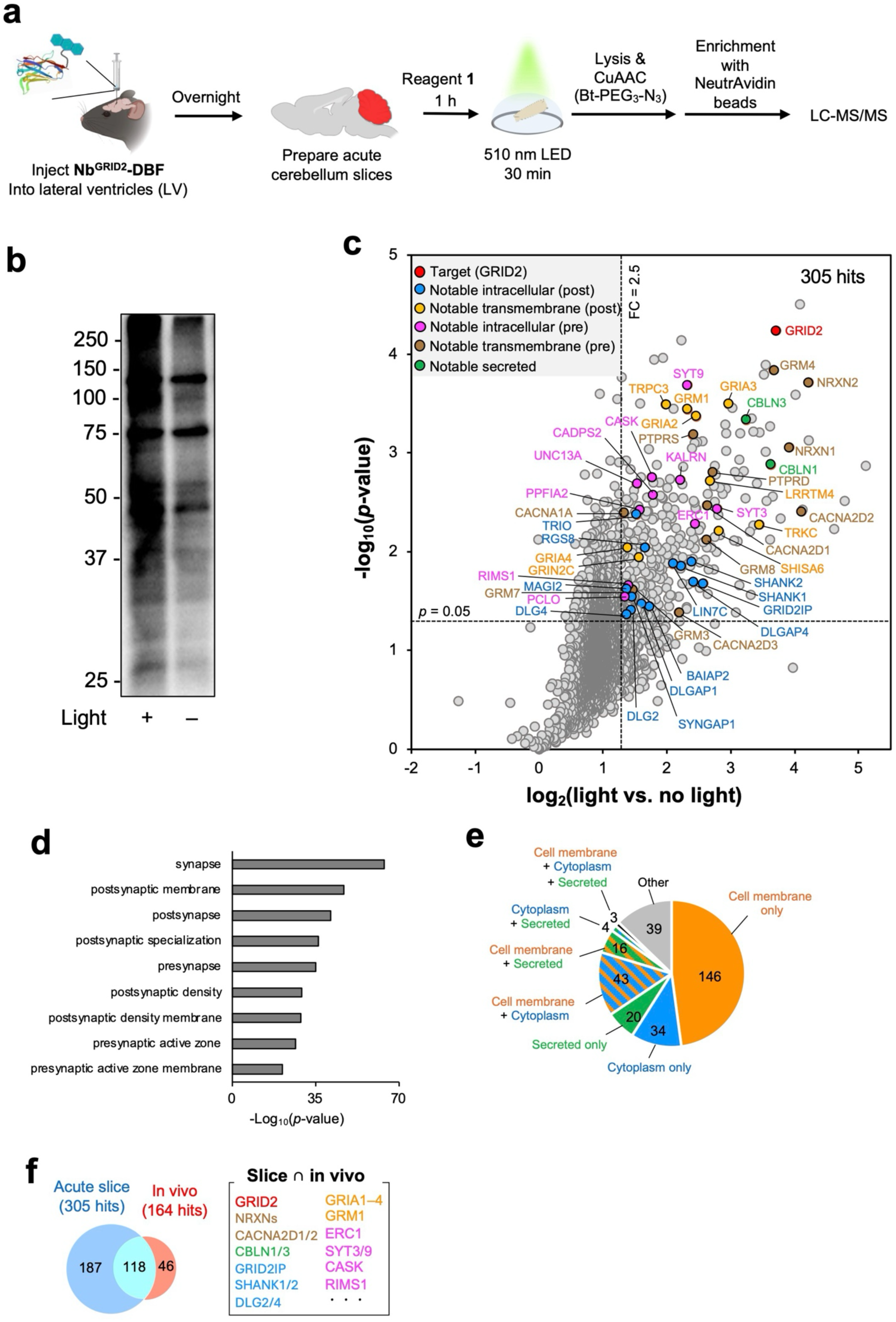
tmPhoxID can be performed in acute brain slices. (**a**) Experimental workflow for tmPhoxID in acute brain slices. (**b**) Streptavidin blot of the proteins labelled by tmPhoxID in acute cerebellum slices. Western blotting was performed after NeutrAvidin enrichment. The data are representative of at least two independent measurements. (**c**) Volcano plot of proteins identified by LC–MSMS. Hit proteins were defined as those with a FC > 2.5 and *p*-value < 0.05 (n = 3). (**d**) Cellular compartment enrichment analysis with Gene Ontology Cellular Component (GOCC). (**e**) Subcellular location by UniProt annotation of the hit proteins. (**f**) Comparison of hit proteins between GRID2-tmPhoxID in brain slices and *in vivo*. The known GRID2 proximal proteins and major constituent proteins of transsynaptic nanocolumns were detected in both contexts.

**Extended Data Fig. 5.**
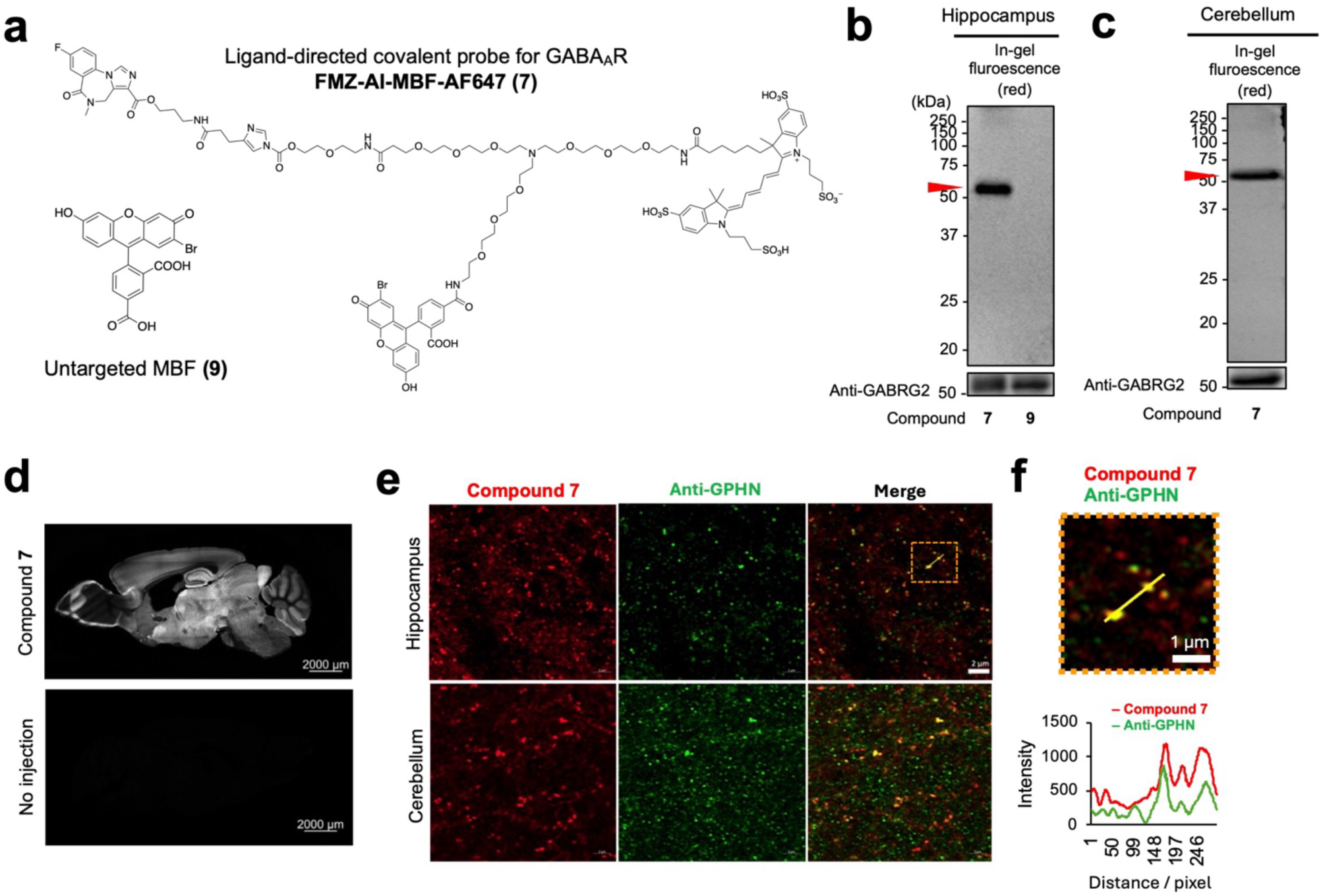
Ligand-directed covalent probe for GABA_A_R. (**a**) Molecular structures of FMZ-AI-MBF-AF647 (**7**) and untargeted control compound (**9**). Compound **7** noncovalently binds to the benzodiazepine-binding site of GABA_A_Rs and covalently transfers the MBF/AF647 moiety to the receptor, driven by the proximity effect. (**b**, **c**) In-gel fluorescence and western blot analysis of (**b**) hippocampus and (**c**) cerebellum tissues from mice injected in the LVs with compound **7** or **9**. The red arrow heads indicate the band corresponding to MBF/AF647-modified GABA_A_Rs. The data are representative of at least two independent measurements. (**d**) Confocal laser scanning microscopy imaging of AF647 fluorescence in whole brain slices prepared from mice injected in the LVs with compound **7**. Scale bar, 2 mm. The data are representative of at least two tissue slices. (**e**) High-resolution confocal images of (upper) hippocampus and (lower) cerebellum prepared from mice injected in the LVs with compound **7**. Scale bar, 2 μm. The anti-gephyrin antibody (Anti-GPHN) was used as a specific inhibitory synapse marker. (**f**) Enlarged view of the orange dashed box and the fluorescence intensity profile along the yellow line. Scale bar, 1 μm.

**Extended Data Fig. 6.**
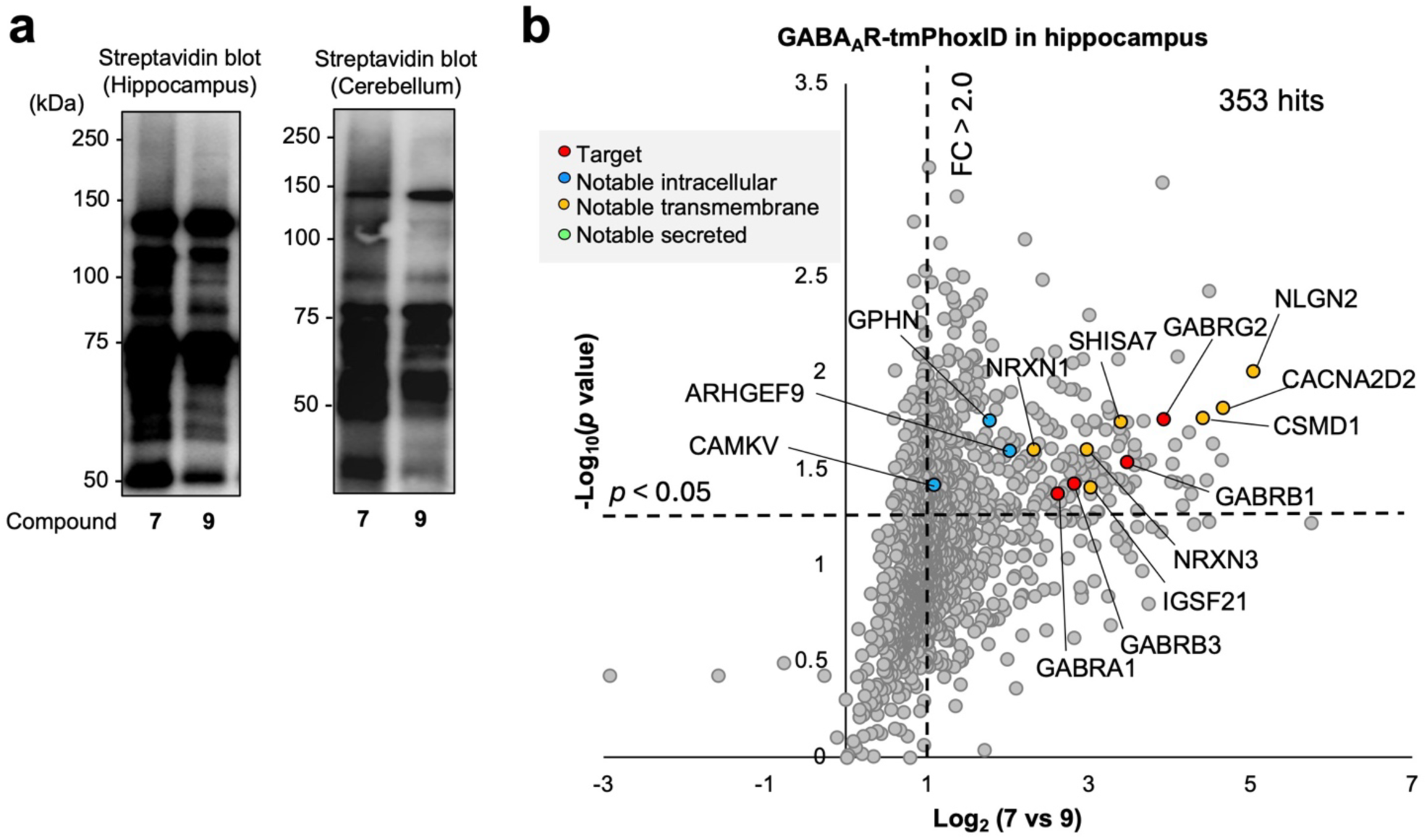
GABA_A_R-tmPhoxID *in vivo*. (**a**) Streptavidin blot of proteins labelled by tmPhoxID in hippocampus and cerebellum in the living mouse brain. Western blotting was performed after NeutrAvidin enrichment. The data is representative of at least two independent measurements. (**b**) Volcano plot of the proteins identified by LC–MSMS. Label-free quantitative mass spectrometry was performed using the untargeted MBF **9** as a control. Hit proteins were defined as those with a FC > 2, *p*-value < 0.05, and unique peptides ≥ 2 (n = 3).

**Extended Data Fig. 7.**
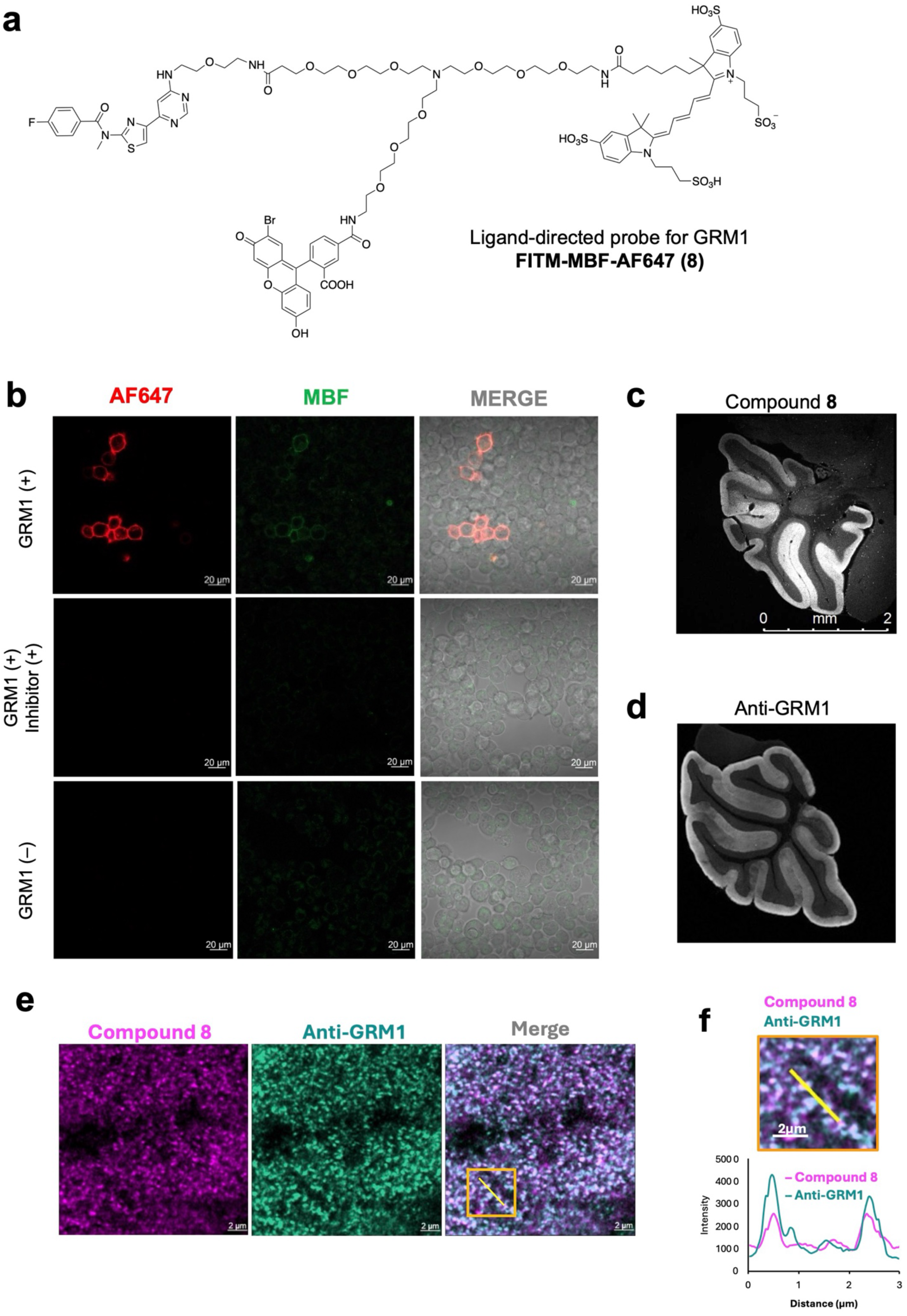
Ligand-directed noncovalent probe for GRM1. (**a**) Molecular structure of FITM-MBF-AF647 (**8**). (**b**) Evaluation of GRM1-selectivity of compound **8** in cultured cells. AF647 fluorescence of compound **8** was strongly detected at the plasma membrane of GRM1-transfected cells, whereas no signal was observed in the presence of the competitive inhibitor FITM or in non-transfected cells. (**c**) Confocal laser scanning microscopy imaging of AF647 fluorescence in cerebellum slices prepared from mice injected in the LVs with **8**. Consistent with the immunostaining images of GRM1 in the mouse cerebellum (**d**), strong fluorescence from compound **8** was detected in the molecular layer, where GRM1 is abundantly expressed. Scale bar, 2 mm. The data are representative of at least two tissue slices. (**e**) High-resolution confocal images of the cerebellar molecular layer stained with compound **8** and anti-GRM1 antibody. The data are representative of at least two tissue slices.

**Extended Data Fig. 8.**
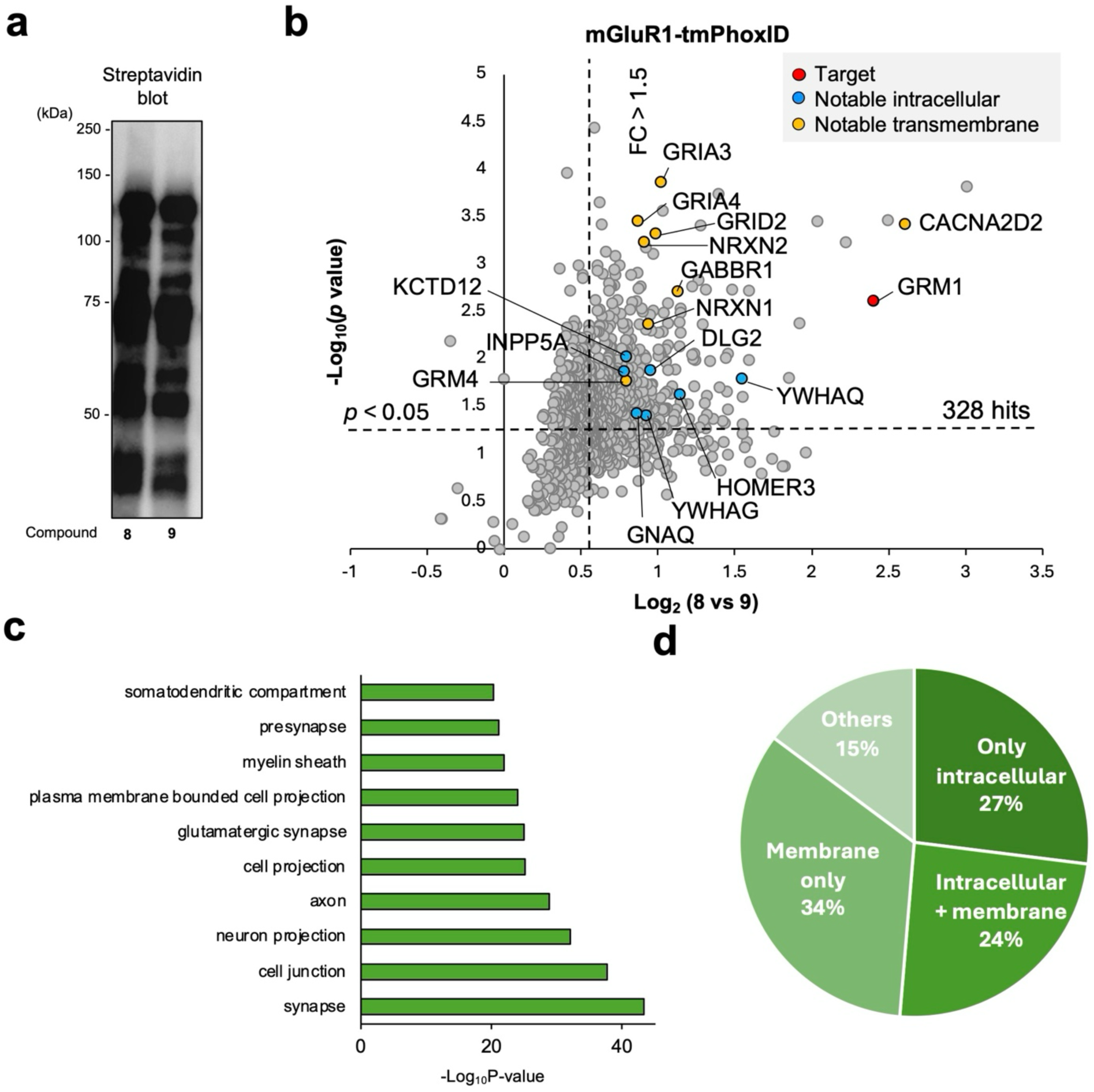
GRM1-tmPhoxID *in vivo*. (**a**) Streptavidin blot of proteins labelled by tmPhoxID using compound **8** or **9** in the cerebellum of living mouse brains. Western blotting was performed after NeutrAvidin enrichment. The data are representative of at least two independent measurements. (b) Volcano plot of proteins identified by LC–MSMS. Label-free quantitative mass spectrometry was performed using untargeted MBF **9** as a control. Hit proteins were defined as those with a FC > 1.7, *p*-value < 0.05, and unique peptides ≥ 7 (n = 3). (**c**) Cellular compartment enrichment analysis with GOCC. (**d**) Subcellular location by UniProt annotation of hit proteins.

**Extended Data Fig. 9.**
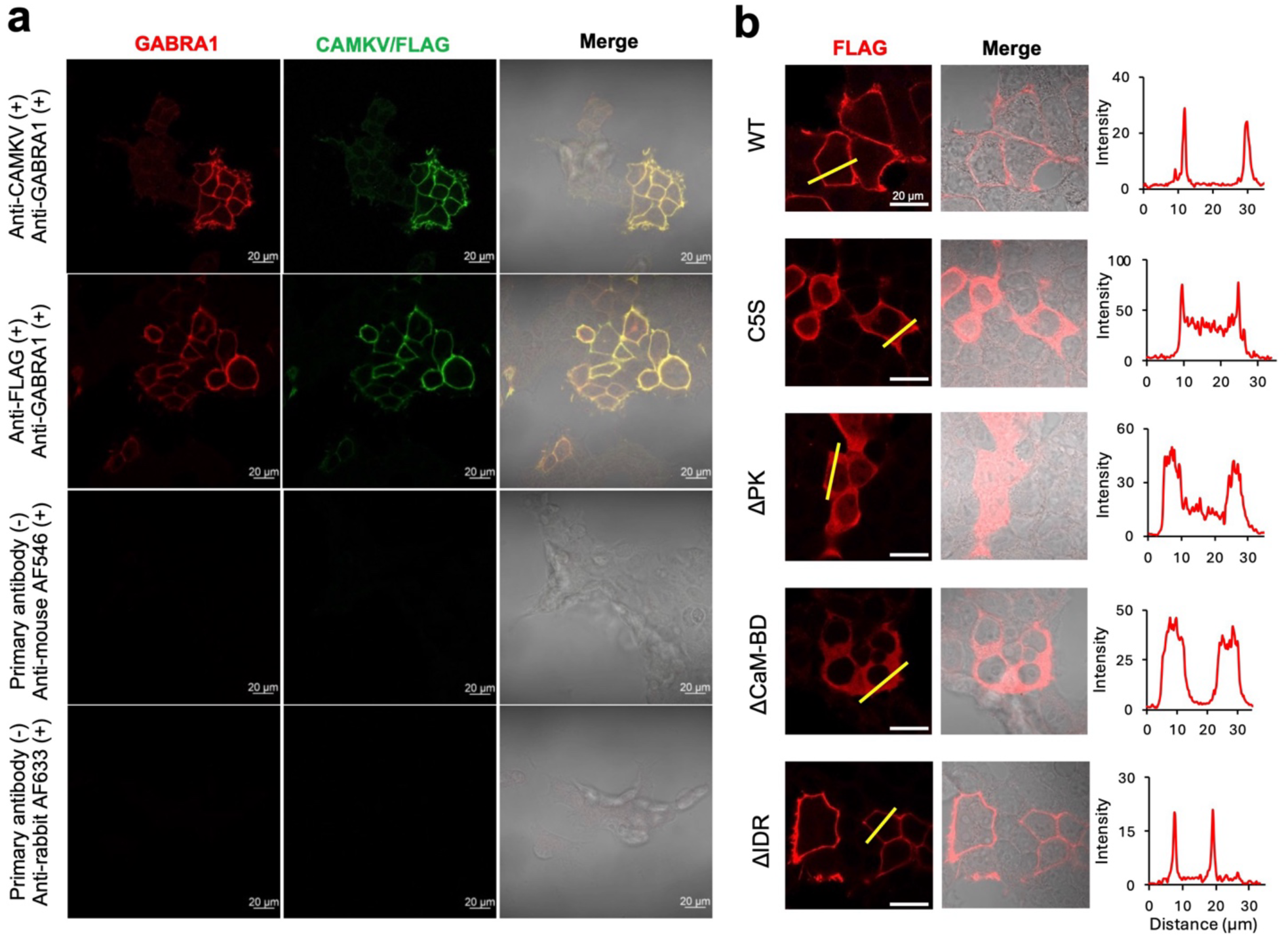
Subcellular localization of CAMKV. (**a**) Immunostaining images of HEK293T cells co-transfected with three GABA_A_R subunits (GABRA1, GABRB3, and GABRG2) and CAMKV-FLAG (WT) indicated that these proteins colocalize at the plasma membrane. GABA_A_R staining was performed using an anti-GABRA1 antibody followed by an anti-rabbit IgG–AF633 conjugate, and CAMKV-FLAG was detected using either an anti-CAMKV or an anti-FLAG antibody followed by an anti-mouse IgG–AF546 conjugate. No fluorescence was detected in controls omitting the transfection (**Supplementary Fig.3**), confirming the specificity of the antibodies used. Scale bar: 20 µm. (**b**) Confocal laser scanning microscopy images showing subcellular localization of CAMKV variants. Whereas the WT and ΔIDR constructs exhibited specific localization to the plasma membrane, the ΔPK and ΔCaM-BD constructs were expressed in the cytosol. The C5S mutant showed enrichment at the plasma membrane but was also distributed in the cytosol. The plot was generated by quantifying the fluorescence intensity along the yellow line. Scale bar: 20 µm.

**Extended Data Fig. 10.**
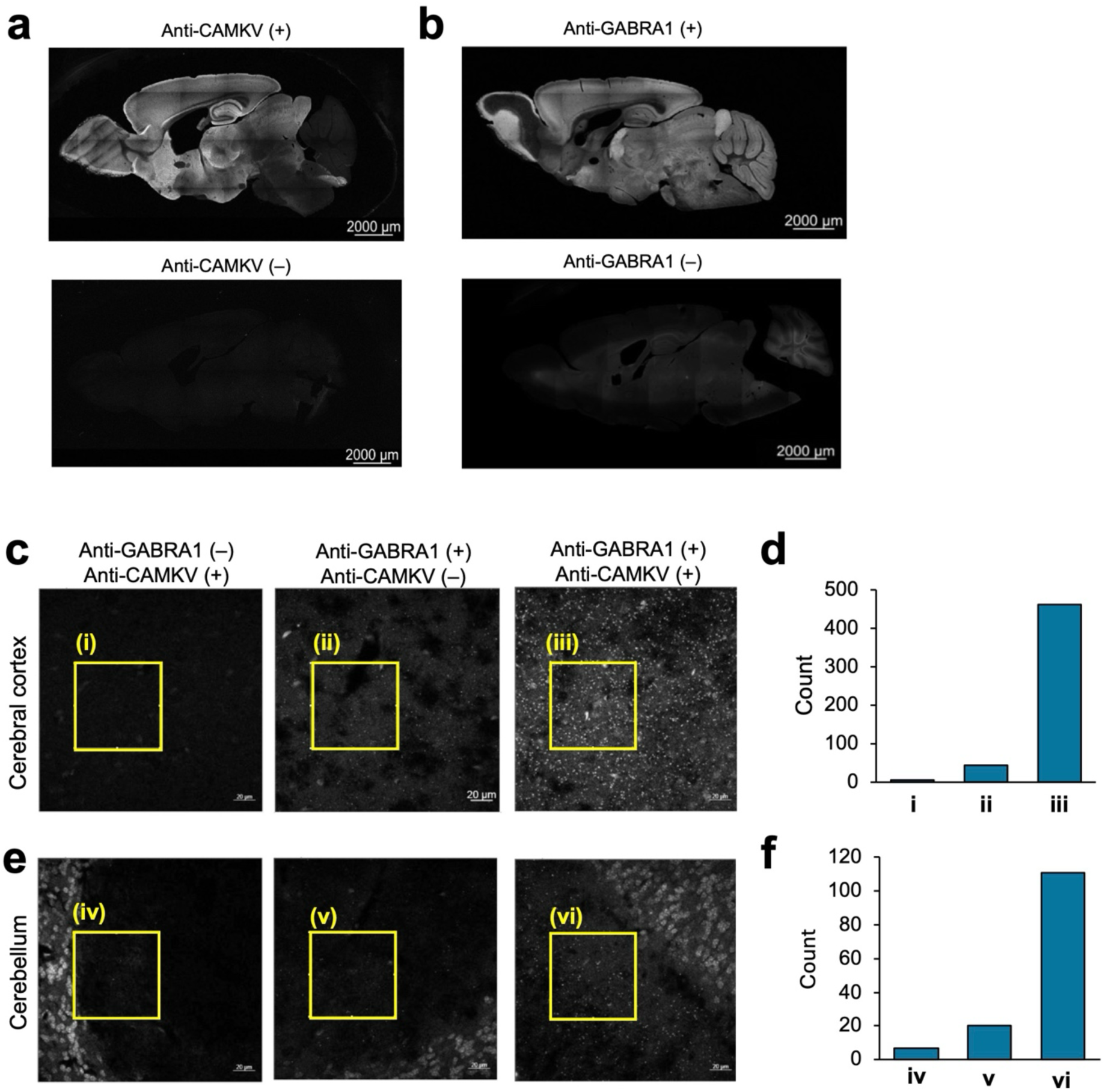
Proximity ligation assay of CAMKV and GABA_A_R in mouse brain tissues. (**a, b**) Confocal laser scanning microscopy imaging of (**a**) CAMKV and (**b**) GABRA1 in whole brain slices stained by anti-CAMKV and anti-GABRA1 antibodies, respectively. Scale bar, 2 mm. The data are representative of at least two tissue slices. (**c**, **e**) PLA assay with anti-CAMKV and anti-GABRA1 antibodies in the mouse (**c**) cerebral cortex and (**e**) cerebellum. (**d**, **f**) Quantification of puncta in the yellow square of panels **c**, **e** (512 × 512 pixels), respectively.

