## Supplementary Information for "Transmembrane PhoxID: Photo-proximity labelling across the plasma membrane"

### Contents

**Supplementary Fig. 1**
**Supplementary Fig. 2**
**Supplementary Fig. 3**

**Supplementary Note | Synthesis methods**

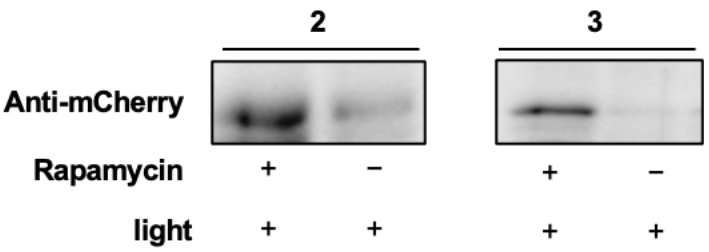


**Supplementary Fig. 1 | tmPhoxID using compounds 2 and 3.** Using the same system as in Figure 2c and d, we confirmed that rapamycin-dependent labeling of cytosolic mCherry–FKBP12 also proceeded when compounds **2** and **3** were used.

| UniprotID | Protein name | Gene name | #unique peptides | Fold change | p value |
|---|---|---|---|---|---|
| Q3UHL1 | CaM kinase-like vesicle-associated protein | Camkv | 2 | 4.44635326 | 0.04117228 |
| Q9WUP7 | Ubiquitin carboxyl-terminal hydrolase isozyme L5 | Uchl5 | 2 | 3.72117237 | 0.0043437 |
| Q9JMB8 | Contactin-6 | Cntn6 | 10 | 3.20014892 | 0.00208924 |
| Q6P1Y8 | Type II inositol 3,4-bisphosphate 4-phosphatase | Inpp4b | 3 | 2.29905492 | 0.01506545 |

**Supplementary Fig. 2 | Four candidate GABA$_A$R-interacting proteins that met our criteria.**

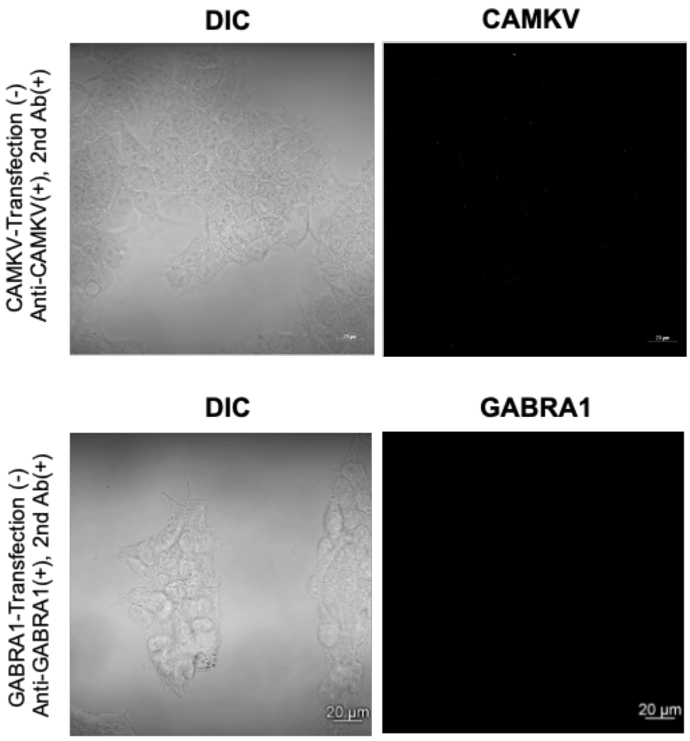


**Supplementary Fig. 3 | Negative control data demonstrating the specificity of anti-CAMKV and anti-GABRA1 used in this study.**

#### Supplementary Note

##### General materials and methods for organic synthesis

All chemical reagents and solvents were obtained from commercial suppliers (Tokyo Chemical Industry (TCI), Sigma-Aldrich, Thermo Fisher Scientific, Fujifilm-Wako Pure Chemical Corporation, Watanabe Chemical Industries, or Kanto Chemical Co., Inc.) and used without further purification. Thin-layer chromatography (TLC) was performed on silica gel 60 F254 or reverse-phase TLC (RP-TLC) plates (60 RP-18 F254s), both precoated on aluminum sheets (Merck). Spots were visualized by fluorescence quenching, fluorescence under 365 nm UV light, or ninhydrin staining. Chromatographic purification was accomplished using a flash column chromatography on silica gel 60 N (neutral, 40–50 μm, Kanto Chemical). $^{1}$H- and $^{13}$C- NMR spectra were recorded in deuterated solvents on a Varian Mercury 400 (400 MHz) spectrometer or JEOL JNM-ECZ600R/S1 (600 MHz), and calibrated to tetramethylsilane (= 0 ppm) or residual solvent peak. Multiplicities are abbreviated as follows: s = singlet, brs= broad singlet, d = doublet, t = triplet, q = quartet, m = multiplet, dd = double doublet. High-resolution mass spectra were measured on an Exactive Plus (Thermo Fisher Scientific) equipped with electron spray ionization (ESI). Reversed-phase HPLC (RP-HPLC) was carried out on a Hitachi Chromaster system equipped with a 5410 UV detector and a 5420 UV-Vis detector.

##### Synthesis

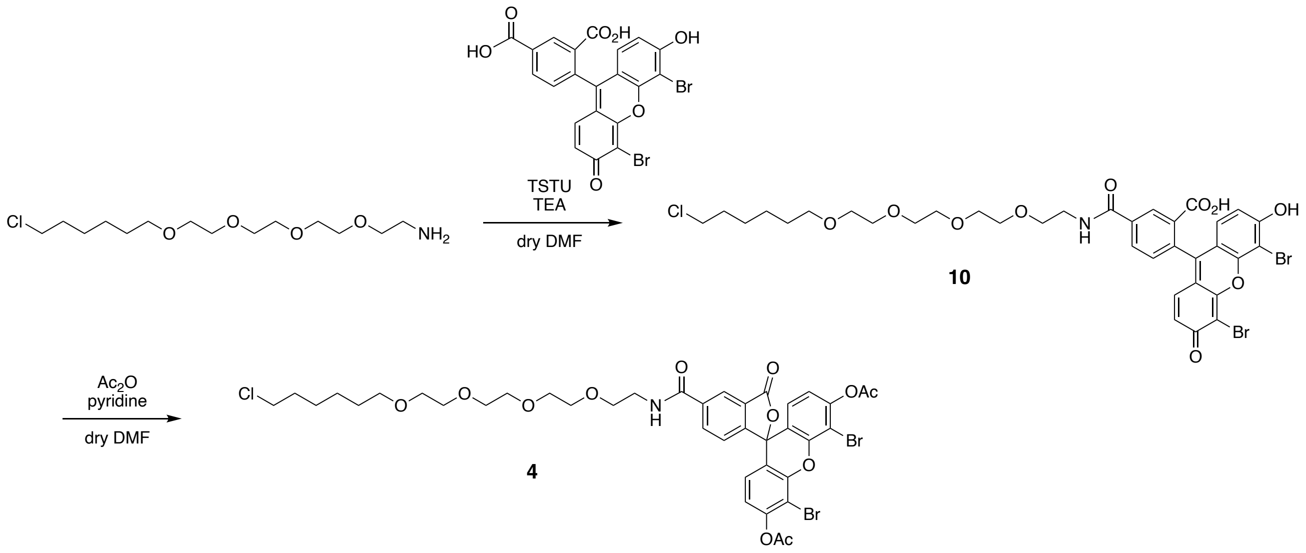


**Scheme 1** Synthetic scheme of compound **4 (AcDBF-HL)**

*Synthesis of* ***10***

To a solution of **DBF-COOH**[S1] (20.0 mg, 37 μmol) in dry DMF, *N,N,N',N'*-Tetramethyl-*O*-(*N*-succinimidyl)uronium Tetrafluoroborate (TSTU) (11.0 mg, 37 μmol) and triethylamine (TEA) (16 μL, 111 μmol) was added at r.t.. After stirring for 2h, a solution of 18-chloro-3,6,9,12-tetraoxaoctadecan-1-amine (12.7 mg, 41 μmol) in dry DMF (0.4 mL) was added to the reaction mixture and stirred overnight at r.t.. After evaporating, the crude product was purified by flash column chromatography (silica, $CHCl_3$ : MeOH = 5:1 + 0.1% TFA), evaporated and dried under vacuum. Target compound **10** was obtained as an orange oil. The crude product was used in the next step without further purification.

*Synthesis of **4***

To a solution of **10** in dry DMF (1 mL), pyridine (58.8 mg, 744 μmol) and $Ac_2O$ (19.4 mg, 190 μmol) was added and stirred at r.t. for 1h. After drying under vacuum, the crude product was diluted with 0.1 M HCl (20 mL) and extracted with ethyl acetate (50 mL). The organic layer was washed with brine, dried over $MgSO_4$ and filtered. The filtrate was concentrated and purified by flash column chromatography (silica, $CHCl_3$ ; MeOH = 60:1) to affordin target impure compound **8**. Obtained product was repurified by flash column chromatography (silica, acetonitrile : hexane = 1:1) to afford pure **8** as colourless oil (2.6 mg, 2.9 μmol, two step 7.8%).

$^1$H-NMR (600 MHz, $CDCl_3$): δ 8.52 (1H, d, *J* = 1.8 Hz ), 8.29 (1H, dd, *J* = 1.8, 8.4 Hz), 7.70 (1H, brd), 7.28 (1H, *J* = 8.4 Hz), 6.93 (2H, d, *J* = 9.0 Hz), 6.81 (2H, d, *J* =9.0 Hz), 3.72–3.65 (12H, m), 3.62–3.60 (2H, m), 3.54–3.53 (2H, m), 3.50 (2H, t, *J* = 6.6 Hz), 3.40 (2H, t, *J* = 6.6 Hz), 2.40 (6H, s), 1.74–1.69 (2H, m), 1.53–1.49 (2H, m), 1.42–1.36 (2H, m), 1.33–1.30 (2H, m).

$^{13}$C-NMR (150 MHz, $CDCl_3$): δ 169.11, 167.98, 167.87, 165.38, 154.31, 150.62, 148.99, 137.49, 135.12, 126.95, 126.11, 124.28, 124.27, 119.56, 117.55, 106.67, 71.22, 70.52, 70.51, 70.47, 70.19, 70.01, 69.78, 45.04, 40.18, 32.48, 29.32, 26.64, 25.57, 25.34, 20.81, 17.60.

HRMS (ESI): Calcd for $[C_{39}H_{42}Br_2ClNO_{12}Na]^+$ 932.0654. Found 932.0658.

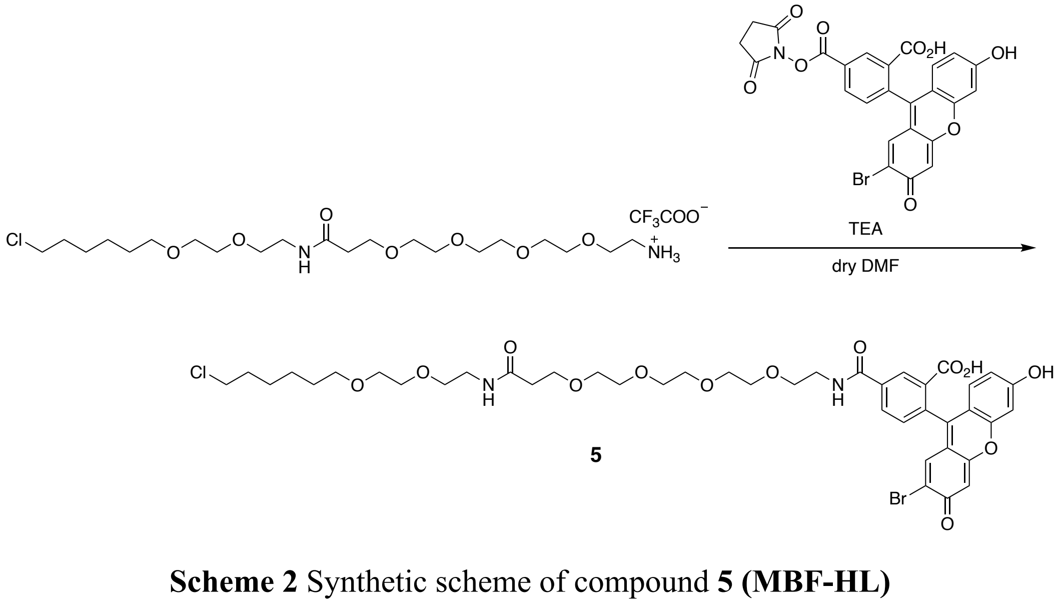


**Scheme 2** Synthetic scheme of compound **5 (MBF-HL)**

*Synthesis of **5***

To a solution of 1-amino-*N*-(2-(2-((6-chlorohexyl)oxy)ethoxy)ethyl)-3,6,9,12-tetraoxapentadecan-15-amide TFA salt (13.3 mg, 22.8 μmol) in dry DMF (500 μL), **MBF-NHS**[S2] (8.4 mg, 15.2 μmol) and TEA (10.5 μL, 76.0 μmol) were added and stirred overnight at r.t.. After evaporating the solvent, the crude product was purified by flash column chromatography on silica gel ($CHCl_3$ : MeOH = 30 : 1) to afford compound **5** (4.33 mg, 4.77 μmol, 31.4%) as yellow oil.

$^1$H-NMR (600 MHz, $CD_3OD$): δ 8.47 (1H, s), 8.24 (1H, dd, $J$ = 7.8, 1.8 Hz), 7.35 (1H, d, $J$ = 8.4 Hz), 6.84 (2H, s), 6.73 (2H, d, $J$ = 1.8 Hz), 6.64 (1H, d, $J$ = 8.4 Hz), 6.58 (1H, dd, $J$ = 1.8, 8.4 Hz), 3.72–3.50 (26H, m), 3.45 (2H, t, $J$ = 6.6 Hz), 3.34 (2H, t, $J$ = 6.4 Hz), 2.42 (2H, t, $J$ = 6.4 Hz), 1.74 (2H, quin, $J$ = 7.0 Hz), 1.56 (2H, quin, $J$ = 7.4 Hz), 1.48–1.29 (4H, m).

$^{13}$C-NMR (150 MHz, $CD_3OD$): δ 174.05, 170.08, 168.56, 162.00, 160.23, 160.01, 154.11, 153.53, 138.31, 135.75, 132.89, 130.34, 128.66, 125.98, 125.45, 117.73, 115.88, 114.26, 113.04, 110.86, 104.51, 103.69, 72.27, 71.66, 71.65, 71.59, 71.50, 71.40, 71.39, 71.32, 71.23, 70.58, 70.47, 68.30, 45.75, 41.32, 40.44, 37.63, 33.79, 30.57, 27.78, 26.52.

HRMS (ESI): Calcd for $[(C_{39}H_{42}Br_2Cl_1N_1O_{12}Na]^+$ 932.0654. Found 932.0662.

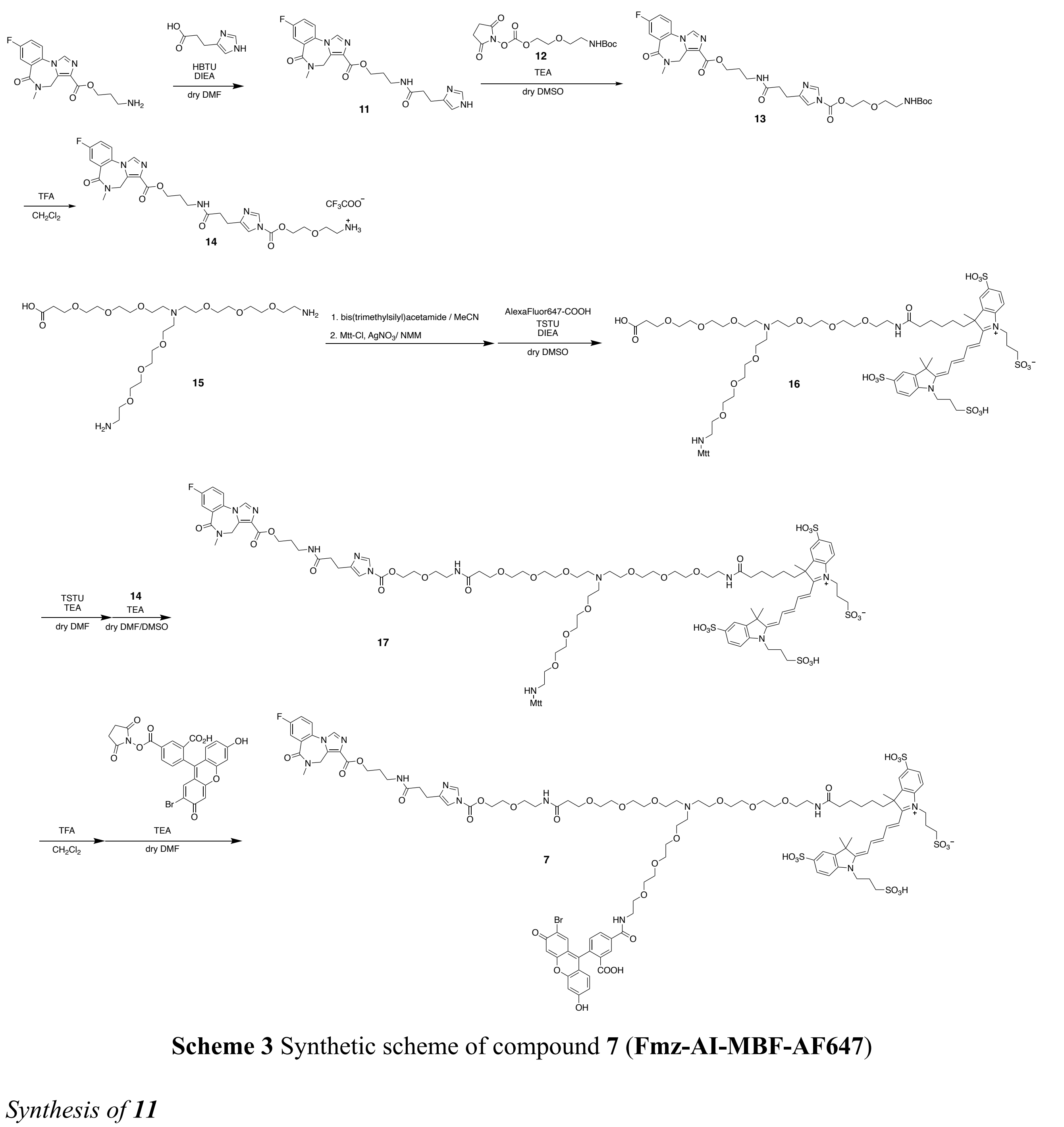

**Scheme 3** Synthetic scheme of compound **7** (**Fmz-AI-MBF-AF647**)

*Synthesis of **11***

To a stirred solution of 4*H*-Imidazo[1,5-*a*][1,4]benzodiazepine-3-carboxylic acid, 8-fluoro-5,6-dihydro-5-methyl-6-oxo-, 3-aminopropyl ester (26.3 mg, 0.079 mmol) in dry DMF (2 mL), 3-(1*H*-imidazol-4-yl)propanoic acid (16.8 mg, 0.12 mmol), 1-[bis(dimethylamino)methylene]-1H-benzotriazolium 3-oxide hexafluorophosphate (HBTU, 45.5 mg, 0.12 mmol), and *N,N*-diisopropylethylamine (DIEA, 150 μL, 0.8 mmol) were added. The reaction mixture was stirred at room temperature under Ar atmosphere for 16 h. After removal of DMF by evaporation, the residue was purified by silica gel chromatography with a linear gradient of 0–40% $CH_3OH/CHCl_3$-1% $NH_3$ (12 CV) to give a compound **11** as a clear film (18.5 mg, 0.041 mmol, 52%).

$^1$H-NMR (400 MHz, $CD_3OD$): δ 7.28 (1H, s), 7.90 (1H, s), 7.76–7.69 (2H, m), 7.55 (1H, s), 7.53–7.49 (1H, m), 6,78 (1H, s), 5.18 (1H, d, *J* = 12.4 Hz), 4.51 (1H, d, *J* = 12.4 Hz), 4.32 (2H, s), 3.34 (2H, t, *J* = 4.2 Hz), 3.22 (3H, s), 2.86 (2H, t, *J* = 7.4 Hz), 2.50 (2H, t, *J* = 7.6 Hz), 1.95 (2H, quin, *J* = 6.5 Hz).

*Synthesis of* ***13***

To a stirring solution of compound **11** (18 mg, 0.040 mmol) in 200 µl of dry DMSO, compound **12** (25 mg, 0.092 mmol) and TEA (64 µL, 0.46 mmol) were added. The reaction mixture was stirred at room temperature for overnight. The crude product was purified by RP-HPLC (AcCN:10 mM triethylammonium acetate (TEAA) = 20 : 80 (5 min), 50 : 50 (30 min)) with a YMC-Pack ODS-A column (5 µm, 250 × 20 mm) at a flow rate of 9.9 mL/min. The target fraction was lyophilized affording compound **13** (15.8 mg, 0.023 mmol, 50%) as a colorless oil.

$^1$H-NMR (400 MHz, $CDCl_3$): δ 8.04 (1H, d, *J* = 0.7 Hz), 7.96, 7.78 (1H, dd, *J* = 4.3, 1.5 Hz), 7.45 (1H, ddd, *J* = 4.4, 2.3, 0.8 Hz), 7.36 (1H, tdd, *J* = 3.6, 1.5, 0.5 Hz), 7.19 (1H, q, *J* = 0.5 Hz), 4.53 – 4.48 (2H, m), 4.39 (2H, d, *J* = 10.0 Hz), 3.78 – 3.74 (2H, m), 3.55 (2H, t, *J* = 2.6 Hz), 3.45, 3.31 (q, *J* = 2.7 Hz), 3.24 (3H, d, *J* = 0.8 Hz), 2.90 (td, *J* = 3.7, 0.5 Hz), 2.61 (t, *J* = 3.6 Hz), 1.96 (dt, *J* = 8.3, 3.0 Hz), 1.43 (9H, s).

MALDI-TOF-MS (+): Calcd for $[C_{32}H_{42}N_7O_9]^+$ $[M\text{-}F\text{+}H]^+$: 668.7275, found: 668.8423

*Synthesis of* ***14***

To a stirring solution of compound **13** (15.8 mg, 0.023 mmol) in $CH_2Cl_2$ (1 mL), TFA (400 µL) was added and the reaction was stirred at room temperature for 45 min. After the completion of the reaction, the solvent was removed under reduced pressure and co-evaporated with toluene (500 µL x 2). The obtained residue of **14** was dried under vacuum for 2 h and was used in the next step of the synthesis without any further purification.

Synthesis of **16**

The commercially available compound **15** (3.8 mg, 6.6 µmol) was dissolved in 300 µL AcCN and bis(trimethylsilyl)acetamide (2.23 mg, 9.98 µmol) was added. The mixture was stirred at room temperature for 40 min. Next, the reaction was diluted by adding 300 µL of ice-cold AcCN, and *N*-methylmorpholine (NMM, 1.3 µL, 11.97 µmol), 4-methyltriphenylmethyl chloride (Mtt-Cl, 3.31 mg, 11.31 µmol) and $AgNO_3$ (2.26 mg, 13.3 µmol) were added to the reaction solution. The mixture was stirred at room temperature for 40 min. The reaction was filtered under vacuum, washed with MeOH (2 x 500 µL), and the filtrate was concentrated under reduced pressure. The obtained white residue (Mtt-protected compound) was used in the next step of the synthesis without any further purification.

In a different flask, the commercially available Alexa Fluor™ 647 carboxylic acid tris (triethylammonium) salt (5 mg, 5.2 µmol), TSTU (1.6 mg, 5.3 µmol), and DIEA (6.9 µL, 7.4 µmol) were dissolved in DMF (300 µL). The mixture was stirred at room temperature in the dark for 2 h. The

reaction mixture was combined with the white residue (Mtt-protected compound)(1.79 mg, 2.62 µmol) and was stirred at room temperature overnight in the dark. The obtained crude product was purified by RP-HPLC (AcCN:10 mM TEAA 10:90 (0 min), 40:60 (30 min)) with a YMC-Pack Pro C18 RS column (5 µm, 250 × 10 mm) at a flow rate of 3.0 mL/min. Target fraction was lyophilized affording compound **16** (1.56 mg, 0.935 µmol, 22%) as a blue film.
MALDI-TOF-MS (-): Calcd for $[C_{61}H_{96}N_5O_{24}S_4]^-$ $[M\text{-}Mtt\text{-}H]^-$: 1411.6905, found: 1411.9230

*Synthesis of **17***

To a stirring solution of compound **16** (1.56 mg, 0.935 µmol) in 300 µL dry DMF, TSTU (0.42 mg, 1.402 µmol) and TEA (3.8 µL, 28.05 µmol) were added. The mixture was stirred at room temperature for 1 h and the completion of the reaction was confirmed by RP-TLC analysis (AcCN : 10mM TEAA = 1 : 2). To the reaction solution, a solution of compound **14** (1.79 mg, 2.62 µmol) in dry DMSO (50 µL) and TEA (3.8 µL, 28.05 µmol) were added. The resulting mixture was stirred at room temperature overnight and the solvent was removed under vacuum. The obtained residue was purified by RP-HPLC (AcCN : 10 mM TEAA = 10 : 90 (0 min), 40 : 60 (30 min)) with a YMC-Pack Pro C18 RS column (5 µm, 250 × 10 mm) at a flow rate of 3.0 mL/min. The target fraction was lyophilized affording target compound **17** (1.09 mg, 0.486 µmol, 54%) as a blue film.
MALDI-TOF-MS (+): Calcd for $[C_{88}H_{129}FN_{12}O_{30}S_4]^{2+}$ $[M\text{-}Mtt+2H]^{2+}$: 1980.2779, found: 1980.7233.

*Synthesis of 7*

Compound **17** (1.09 mg, 0.486 µmol) was dissolved in 500 µL $CH_2Cl_2$ with 1% TFA. The solution was stirred at room temperature for 30 min. The completion of the reaction was confirmed by RP-TLC analysis (AcCN : 10 mM TEAA = 1 : 2), and the solvent was co-evaporated with toluene (500 µL x 2). To a solution of the obtained residue in dry DMF (100 µL), **MBF-NHS** (0.493 mg, 0.729 µmol,) and TEA (0.337 µL, 2.43 µmol) were added. The reaction was stirred at room temperature overnight. RP-TLC analysis (AcCN : 10 mM TEAA = 1 : 2) revealed practically full conversion, and the solvent was removed under vacuum. The obtained residue was purified by RP-HPLC (AcCN : 10 mM TEAA = 10 : 90 (0 min), 40 : 60 (30 min)) with a YMC-Pack Pro C18 RS column (5 µm, 250 × 10 mm) at a flow rate of 3.0 mL/min. The target fraction was lyophilized affording target compound **7** (0.456 mg, 0.198 µmol, 40%) as a blue/green film.
HRMS (ESI): Calcd for $(C_{109}H_{132}BrFN_{12}O_{36}S_4Na_2)^{2-}$ $[M–2H]^{2-}$: 1228.8378, found: 1228.8389.

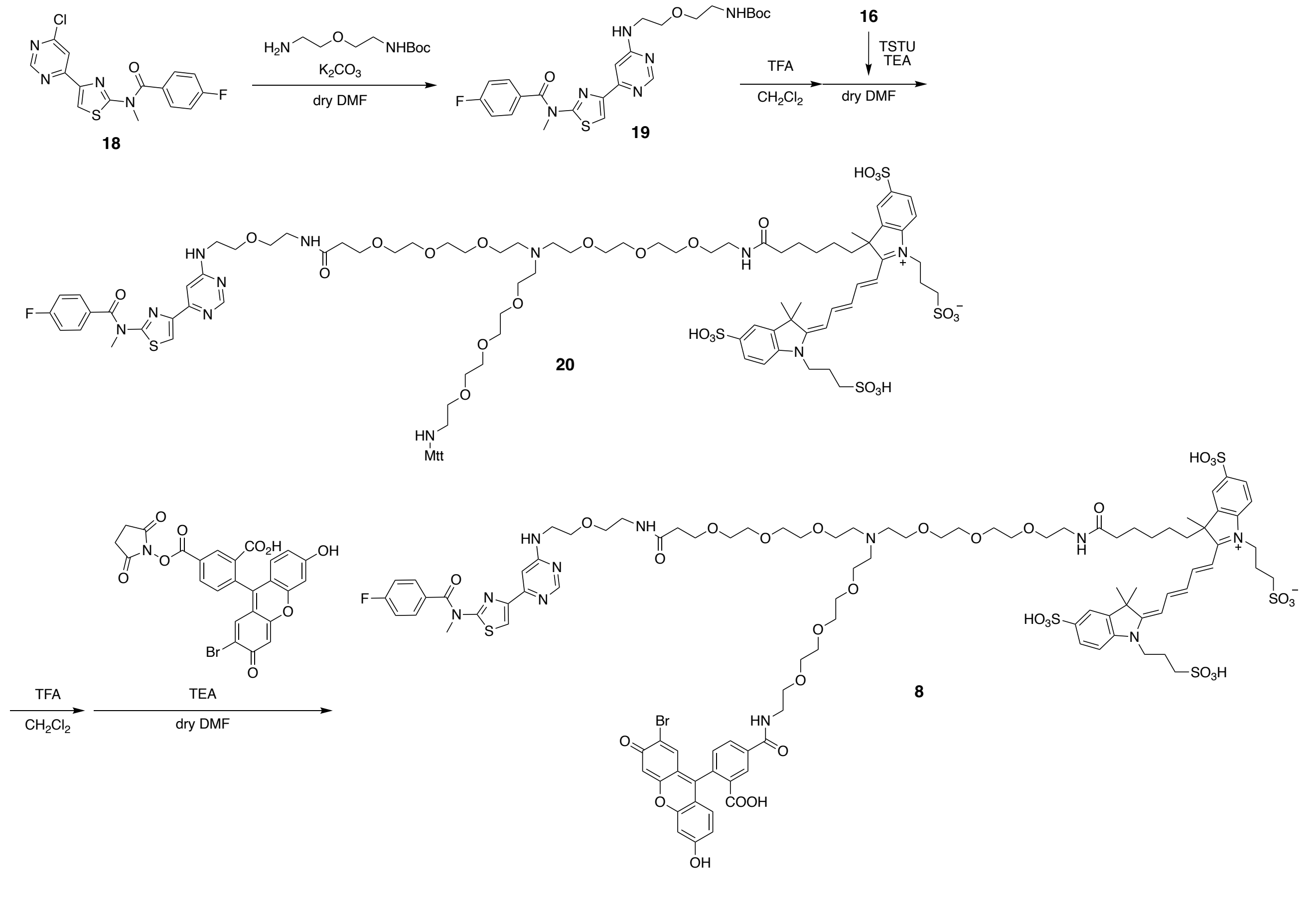

**Scheme 4** Synthetic scheme of compound **8** (**FITM-MBF-AF647**)

*Synthesis of **19***

To a stirring solution of compound **18**[S3] (100 mg, 0.29 mmol) in dry DMF (400 µL), *tert*-butyl (2-(2-aminoethoxy)ethyl)carbamate (86.6 µL, 0.43 mmol) and $K_2CO_3$ (200 mg, 1.54 mmol) were added. The reaction mixture was stirred at room temperature overnight and the completion of the reaction was confirmed by TLC analysis ($CHCl_3$ : MeOH = 10 : 1). The solvent was removed under vacuum, and the obtained residue was purified by flash column chromatography on silica gel ($CHCl_3$ : 1% MeOH) affording target compound **19** (69.5 mg, 47%) as a colorless oil.

$^1$H-NMR (400 MHz, $CDCl_3$): δ 8.57 (d, $J$ = 1.1 Hz, 1H), 7.99 (s, 1H), 7.61 – 7.57 (m, 2H), 7.20 – 7.15 (m, 2H), 7.11 (d, $J$ = 1.2 Hz, 1H), 3.72 (s, 3H), 3.67 – 3.58 (m, 5H), 3.53 (t, $J$ = 5.2 Hz, 2H), 3.32 (t, $J$ = 5.6 Hz, 3H), 1.42 (s, 10H).

*Synthesis of **20***

To a stirred solution of compound **19** (0.7 mg, 1.34 µmol) in $CH_2Cl_2$ (500 µL), TFA (100 µL) was added. The mixture was stirred for 30 min, and the solvent was co-evaporated with toluene (3 x 500 µL). The deprotected product was used without further purification. In a different flask, compound **16** (0.48 mg, 0.803 µmol) was dissolved in 300 µL DMF and mixed with TSTU (0.42 mg, 1.402 µmol) and TEA (3.8 µL, 28.05 µmol). The mixture was stirred at room temperature for 1 h and the completion

of the reaction was confirmed by RP-TLC analysis (AcCN:10mM TEAA = 1 : 2). The obtained reaction solution was mixed with the deprotected product of compound **19** and was stirred at room temperature overnight. RP-TLC analysis (AcCN : 10 mM TEAA = 2 : 1) revealed full conversion of the starting material. The solvent was removed under vacuum and the obtained residue was purified by RP-HPLC (AcCN : 10mM TEAA = 20 : 80 (0 min), 65 : 35 (30 min)) with a YMC-Pack Pro C18 RS column (5 μm, 250 × 10 mm) at a flow rate of 3.0 mL/min. Target fraction was lyophilized affording target molecule **20** (0.256 mg, 0.125 μmol, 26%) as a blue film.
HRMS (ESI): Calcd for $(C_{109}H_{132}BrFN_{12}O_{36}S_4Na_2)^{2-}$ [M-H-Mtt]: 1811.1595, found: 1811.0432.

*Synthesis of **8***

Compound **20** (0.256 mg, 0.125 μmol) was dissolved in 500 μL $CH_2Cl_2$ with 1% TFA. The solution was stirred at room temperature for 30 min. After confirming the completion of the reaction by RP-TLC analysis (AcCN : 10 mM TEAA = 1 : 2), the solvent was co-evaporated with toluene (2 x 500 μL). To the obtained residue, a solution of MBF-NHS (0.102 mg, 0.186μmol) in dry DMSO (300 μL) and DIEA (3 μL) was added, and the mixture was stirred for 4 h at room temperature. The completion of the reaction was confirmed by RP-TLC (AcCN : 10 mM TEAA = 1 : 2) and the reaction mixture was purified by RP-HPLC (AcCN : 10 mM TEAA = 10 : 90 (0 min), 40 : 60 (30 min)) with a YMC-Pack Pro C18 RS column (5 μm, 250 × 10 mm) at a flow rate of 3.0 mL/min. Target fraction was lyophilized affording target compound **8** (0.120 mg, 0.054 μmol, 43%) as a greenish blue film.
HRMS (ESI): Calcd for $[C_{101}H_{121}BrFN_{11}O_{31}S_5Na_2]^{2-}$: 1143.7904, found: 1143.7989.
